# Song patterns support species status for some, but possibly not all, island populations of House Wren (*Troglodytes spp.*) in the Lesser Antilles

**DOI:** 10.64898/2026.01.15.699782

**Authors:** Drew Rendall

## Abstract

Island populations are special for the study of evolutionary processes and can be a zone of incipient speciation. Recently, several island populations of House Wren in the Lesser Antilles (Dominica, St Lucia, St Vincent, and Grenada), formerly recognized as subspecies of the continental form, were reclassified as distinct species. However, much of the supporting data was fragmentary in its sampling of the different islands or equivocal in the patterns observed. Because song is a core element of mate recognition and choice, and can therefore be a key character in species identification, I report here the first detailed characterization and analysis of song for House Wren on all of the islands of the Lesser Antilles where they remain, including Trinidad and Tobago; and compare song patterns across the different islands as well as to several continental populations.

Results show that song is broadly similar across all of the islands and to continental populations in high-level features of its structure, organization and delivery but is discriminably different among many of them in its more detailed features. The latter differences are consistent with the recent species splits, with the possible exception of Grenada. They also support retention of House Wren on Trinidad and Tobago as subspecies of the continental form. Results also point to the possibility of a central American origin for some of the islands and a south American origin for others, yielding a trait mosaic where islands that putatively share the same geographic origins, and are therefore presumably genetically closest, are not the most similar in patterns of song (or plumage). This pattern would therefore entail multiple intriguing instances of convergent evolutionary divergence among them that warrants further detailed study.

**Lay Summary:**

- I provide the first comprehensive analysis and comparison of song patterns of House Wrens for all of the islands of the Lesser Antilles where they remain, some of which are at risk of extirpation, or even extinction if they represent distinct species.
- I use the patterns to interpret the recent taxonomic reclassification of many of these island populations as distinct species.
- In their general structure, organization and delivery male song is similar across all of the islands and follows patterns common to contintental forms of House Wren distributed broadly across North, Central and South America.
- Songs of the different islands are, however, discriminably different in their more detailed features and these differences are consistent with most, but possibly not all, of the recent species splits.
- For the island populations recently reclassified as different species, the distinctiveness of male song is greatest in Dominica and St Vincent and to a lesser extent also St Lucia, and least distinctive in Grenada. Song in Trinidad and Tobago is not substantively different from populations in mainland South America which supports retaining these two island populations as subspecies of the closest continental forms.
- Song patterns also point to different possible continental sources for some of the island populations: a source in Central America for Dominica and St Lucia; and a source in South America for the rest. If true, this creates multiple instances of convergent evolutionary divergence in trait patterns across the various islands which merits further study.

---

Island populations are special for evolution both because their physical isolation restricts gene flow, which allows accumulation of even neutral changes, and because their often distinct ecologies can also yield adaptive diversification. As a result, island populations may differentiate relatively rapidly and therefore can be zones of incipient speciation. Darwin’s finches of the Galapagos Islands are a classic example: a relatively rapid radiation of different finch species on different islands that vary relatively subtly in plumage and overall size, as well as in beak size and shape based on variation in foraging niche.

One challenge with definitively establishing species status for island populations – or any population – is knowing how much differentiation is enough because, although there are many proposals, there is no definitive guide to how much variation in typical taxonomic characters, including genes, is required (reviewed in Sukumaran and Knowles 2017; Winker 2021). And the insularity of island populations complicates matters because it effectively precludes interbreeding even if it might occur otherwise. Here song is often given some priority in taxonomic reviews given its role in mate attraction, recognition and mate choice (Remsen 2005; see also Alström and Ranft 2003), and there are many examples of cryptic speciation revealed largely or only on the basis of differences in song patterns (e.g., Irwin 2000; Valderrama et al. 2007; Toews and Irwin 2008). Notably, Podos and colleagues (Podos et al. 2004; Huber & Podos 2006; Podos 2010) documented song variation within and among several species of Darwin’s finches traceable to differences in beak size and shape, illustrating how divergence in feeding niches can have knock-on effects for factors more directly tied to species recognition and status (see also Grant and Grant 1996).

The House Wren (*Troglydytes spp.*) is another species complex well suited to contributing to these evolutionary questions. House Wrens are widely distributed across North, Central and South America. Indeed, they have one of the widest latitudinal distributions of any songbird in the western hemisphere, breeding from approximately 55^0^ north in Canada to 55^0^ south at the southern tip of South America. Across this vast range, house wrens inhabit a diversity of environments and include virtually continuously distributed continental populations but also several island populations in Mexico, Central and South America, and the Lesser Antilles. Hence, it is a taxon that is well suited to examining how variable environments and ecologies affect migration, social and mating systems, and life-history traits, and there is a large and important literature on many of these topics (see taxon reviews by Fernandez et al. 2024; Johnson 2024).

With widely distributed continental forms but also a number of insular island forms, it is a taxon that is also especially well-suited to testing how continuity and discontinuity in distribution can affect population genetic structuring and speciation. This latter important evolutionary issue has long been contested for house wrens, as the classification and species status of populations in this taxon has been complicated and regularly debated. At times, the continental populations have been split into three different species: *T. aedon* (North America), *T. brunneicolis* (northern Mexico), and *T. musculus* (Central and South America), while at other times they have been lumped into a single super-species (*T. aedon*), with more than 30 different subspecies (see for example, Brumfield and Caparella 1996; Brewer 2001; Kroodsma and Brewer 2005; Dickinson and Christidis 2014).

Until recently, the latter framework of a single super-species was formally accepted but not without continuing calls to recognize species status for one or more of the continental forms, as well as several of the island forms. And, indeed, a very recent re-evaluation (Chesser et al. 2024) restored two of the formerly recognized continental species (the northern House Wren: *T. aedon*; and the southern House Wren: *T. musculus*) and elevated to species status three of the formerly recognized island subspecies off the coast of Mexico (Isla Cozumel: *T. beani*; Isla Socorro: *T. sissoni*; Isla Clarion: *T. tanneri*) as well as all of the island subspecies in the Lesser Antilles (Dominica: *T. martenicensis*; St Lucia: *T. mesoleucus*; St Vincent: *T. musicus*; Grenada: *T. grenadensis*) with the exception of those closest to the mainland on Trinidad (*T.m. clarus*) and Tobago (*T.m. tobagensis*).

These recent taxonomic changes were based on a mix of genetic, morphological, plumage, and song characteristics (Chesser et al. 2024). The data supporting species status for the two continental forms and the island forms near Mexico appear clear and compelling. The genetic data involved both mitochondrial and nuclear DNA for a large sample of house wrens across their broad continental distribution, as well samples from all of the island populations of Mexico (Klicka et al 2023); while the song data involved a similarly comprehensive sample for continental populations as well as the island forms in Mexico, including experimental responses to song playbacks for the latter island populations (Sosa-López and Mennill 2014a; Sosa-López et al. 2016; see also Sosa-López and Mennill 2014b,c).

However, the data in support of species status for each of the island populations in the Lesser Antilles are less clear and complete. For example, the genetic sample for the Lesser Antilles group did not include either St. Lucia or Tobago; and it involved only mitochondrial DNA and not nuclear DNA (Klicka et al 2023). The mtDNA results suggested distinct genetic structuring for the island forms that were sampled, as would be expected given their insularity, and also pointed to an intriguing possible difference in the source of colonization of the different islands either from mainland South America or Central America. However, the authors themselves ultimately favored using their nuclear DNA data to resolve taxonomic relationships in the House Wren complex because the mtDNA produced some odd instances of paraphyly that were inconsistent with traditional taxonomy and the general geography (Klicka et al. 2023). Thus, the authors concluded that, “*The greater consistency of the RADseq [nuclear DNA] results and traditional taxonomy and geography suggests it is the more trustworthy phylogenetic pattern … the RADseq data provide the most appropriate basis for classification and understanding House Wren evolution*…”. Consequently, these authors recommended recognizing separate species status for northern and southern House Wren, as well as three of the island forms from Mexico, but they specifically did not make any recommendations concerning the island populations in the Lesser Antilles.

The song data for the Lesser Antilles are also limited. Although their songs are often popularly described as distinctive, there has been no systematic analysis and comparison of song across the various islands. The most relevant and comprehensive comparative work cited by the classification committee is the study by Sosa- López and Mennill (2014). While comprehensive in every other respect, that study was limited in its sampling for the Lesser Antilles: it included only two songs from Dominica, one of which was noted as possibly being from St Vincent; there were no song samples included for any of the other islands. Unfortunately, there are few other formal sources that speak compellingly to the matter of song differences among the islands. There are only a handful of short reports, often with a different primary focus, that offer only subjective, impressionistic descriptions of one or a few songs from a single island. For example, Gilardi and John (1998), also cited by the classification committee, performed an important census and study of the general behavior and breeding biology of house wrens in St Lucia, which are seriously threatened on that island. Their paper is most valuable in other respects but provides only a single spectrogram of one song noting that songs are “highly variable”. Similarly, there is an informal report by Barlow (1978) for house wrens on Gaudeloupe, where the birds are now believed extinct, which provides a spectrogram of one song from that island compared to a recording made of a house wren in Florida (which is not typically part of the species’ breeding range) noting that “to my ear, the song of the Gaudeloupe House Wren is the louder, richer and more melodic of the two”. More recently, Cyr et al. (2021) conducted a much more systematic study of song for Grenada; however, that study was focused specifically on the issue of anthropogenic influences on song for birds living in urban (disturbed) versus rural (undisturbed) locations only on the island of Grenada and involved no data or comparisons to song for any other islands in the Lesser Antilles. Hence, to date there simply is no comprehensive study and comparison of song patterns available for the island group of house wrens in the Lesser Antilles.

There is obvious plumage variation among the island forms, as also noted in the recent reclassification, and illustrated in Figure 1. However, here again, the patterns have been described only historically and in impressionistic terms (Oberholser 1904); they have not been studied more recently or systematically, and the patterns of variation are not entirely straightforward. Thus, birds on the islands closest to the mainland in Trinidad and Tobago are most similar to continental forms, with birds in Tobago potentially moderately distinctive in having a more uniform (bleached) white chest and belly, which in continental forms is often more greyish, cream or buff-colored with a pale russet to rose-colored wash on the flanks. The four more remote islands of the Lesser Antilles show a “leap-frog” pattern of plumage variation (sensu Remsen 1984). Birds on the most southern and most northern of the islands (Grenada and Dominica) are most similar and are much darker overall compared to continental forms, with a more uniformly rufescent color all over and therefore lacking the grey-white underbelly common to continental forms. And the two middle island forms (St Vincent and St Lucia) are also most alike and differ from both Grenada and Dominica and from continental forms in showing a more bounded pattern of white ventrum and pale cheeks (auriculars) combined with chestnut-to-russet colored wings and back, and a comparatively heavy and conspicuous white eye-stripe. The latter eye-stripe is present but variable and generally less prominently developed in the other island populations as well as in most continental forms. Hence, while the four more isolated island populations of the Lesser Antilles recently elevated to species status are genuinely distinctively patterned relative to continental forms, the pattern of plumage variation among them does not suggest four distinctly different color morphs in support of four different species, nor clear clinal variation across them, but rather two clusters with a disjunct distribution. Ultimately, while plumage patterns can often be a useful taxonomic character, they are also famously complicated, inherently subject to environmental variation, and therefore often differing within species with broad distributions (see below).

**Figure 1:**
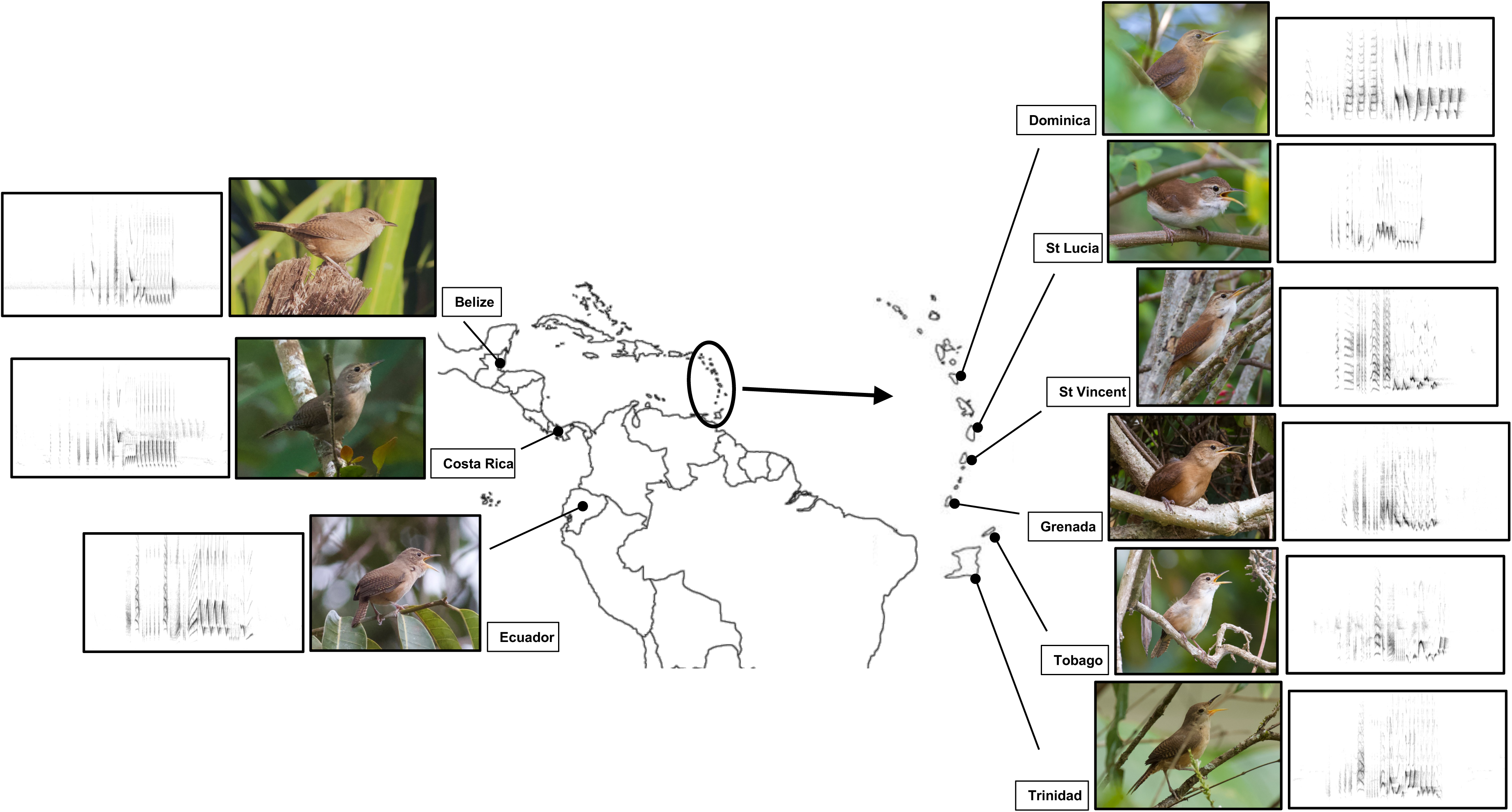
Map of study locations with spectrograms of representative songs and photographs of plumage patterns for each population. Note the ‘leap-frog’ pattern of plumage similarity and difference for the four core islands of the Lesser Antilles that were recently reclassified as separate species, where plumage is similar between Dominica and Grenada at the northern and southern extremes of this group, and also similar between St Lucia and St Vincent in the middle.

Finally, there is well-documented morphological variation in the island populations of the Lesser Antilles (Sosa-López and Mennill 2014; Wetten 2021). In a particularly comprehensive recent study, Wetten (2021) confirmed that the island populations of house wrens are larger in most standard measures of body and beak size compared to mainland populations, and that these measures also differ among some, but not all, of the core islands of the Lesser Antilles (not including Trinidad or Tobago). However, the pattern of larger body and beak size was not unique to the island populations of the Lesser Antilles but applied also to the island populations in Mexico. It was thus interpreted to be an instance of the broader phenomenon of insular gigantism, where small-bodied species tend to get larger on islands, compared to mainland counterparts, as a function of reduced species diversity and attendant ecological release and the adoption of a more generalist niche (Grant 1965; Cox and Ricklefs 1977; Clegg and Owens 2002).

Hence, as with plumage, morphology is an inherently plastic trait subject to considerable environmental variation, even within species; and both characters can thus facilitate but sometimes also complicate species identification. The complications are well illustrated by the example of the North American Fox Sparrow (*Passarella iliaca*) which parallels closely the situation for House Wren. This taxon, widely distributed across North America, has also been a focus of taxonomic debate. It is currently considered a single species, with many subspecies, organized into four broad types (AOU 1998). These four types were once regarded as different species based on variation in body size, beak size and shape, and plumage patterns (Bailey 1902; Swarth 1920, Linsdale 1928), although there is also considerable variation in these traits within each of the types. More recent mtDNA analyses have reported genetic structuring between the four types but with non-trivial amounts of introgression (hybridization) between them, and the morphs that are most similar in size and plumage are not those most similar genetically, indicating that morphology and plumage in this taxon are environmentally plastic and changing independently of gene flow (Zink 1994; Zink and Blackwell 1996; Zink & Weckstein 2003; Zink 2008; for additional details and review see Weckstein et al. 2020). Avian taxonomy includes many similar examples where genetic, size and plumage characteristics do not align neatly (e.g., Yellow-rumped warbler - *Setophaga coronata*: see Hunt & Flashpohler 2020; Northern Flicker – *Colaptes auratus*: see Wiebe & Moore 2024) and where there are therefore no easy taxonomic solutions, only the opportunity that additional data might contribute further insight.

To help address these taxonomic and evolutionary issues for the House Wren group and contribute to the broader enterprise for other taxa, I provide here a first comprehensive characterization and quantitative analysis of song patterns across all of the islands in the Lesser Antilles where house wrens remain. To ground the island comparisons, I include several continental populations in the analysis and compare the complete set to comprehensive descriptions of song reported previously for both northern and southern House Wren (e.g., Kroodsma 1977; Tubaro 1990; Rendall and Kaluthota 2013; dos Santos et al. 2016; DiSciullo et al. 2023).

## METHODS

### Populations Studied

House wrens in the Lesser Antilles were studied twice over a ten-year period, first in 2015 and again in 2024. On both occasions, birds were sampled and recorded on all of the core islands where they remain, which are from north-to-south and in order of increasing proximity to the mainland: Dominica, St Lucia, St Vincent and Grenada. Trinidad and Tobago, which are closest to Grenada and a bit further south, were also deliberately included in this study because they too are part of the Lesser Antilles island chain and because they are also comparatively close to the south American mainland, with correspondingly higher avian species diversity, confirming greater historical and likely also continuing connectivity (i.e., gene flow) with continental populations. Hence, the islands of Trinidad and Tobago represent a natural immediate outgroup from which to root comparisons of song patterns for the other core islands of the Lesser Antilles. Birds on Trinidad and Tobago are also not currently regarded as distinct species but rather as subspecies of the primary south American form (*T. musculus*): the birds in Trinidad belong to the same subspecies as the closest mainland form (*T. m. clarus*); while the birds in Tobago are classified as their own subspecies (*T. m. tobagensis*).

To further root the analysis of island song patterns with continental out-group comparisons, I include several mainland populations from Ecuador, studied in 2022, that are grouped in the same subspecies (*T. m. albicans/clarus*) as birds from Trinidad. I also include several populations from Belize and Costa Rica (hereafter referred to as BCR), studied in 2018 and 2016 respectively, representing additional subspecies of the south American continental form (*T. m. intermedius* and *T. m. inquietus*; see Table 1 and Figure 1 for sample locations). In addition to helping to ground the comparisons of the song patterns of the various island populations, this set of mainland comparisons also facilitates a test of the intriguing possibility raised by the genetic analysis of Klicka et al. (2023) that there might be a different historical source for colonization of the different islands in the Lesser Antilles.

**Table 1.** Summary of song sample.

| Population | Species/subspecies | Dates | Males | Songs | Latitude | Longitude | Elevation (m) |
| --- | --- | --- | --- | --- | --- | --- | --- |
| Belize-Belmopan | *T. m. intermedius* | Mar 7 - Mar 9, 2018 | 6 | 8 | 17.22 | -88.85 | 72 |
| CostaRica-LaSuiza | *T. m. intermedius* | Apr 10 - Apr 14, 2016 | 2 | 8 | 9.85 | -83.61 | 960 |
| CostaRica-Orosi | *T. m. intermedius* | Apr 5 - Apr 9, 2016 | 7 | 16 | 9.78 | -83.84 | 1,250 |
| CostaRica-LasCruces | *T. m. inquietus* | Apr 15 - Apr 19, 2016 | 4 | 16 | 8.78 | -82.95 | 1,218 |
| Dominica-Rosalie | *T. martinicensis* | Apr 21 - Apr 26, 2015 | 6 | 30 | 15.37 | -61.26 | 29 |
| Dominica-Portsmouth |  | Apr 5 - Apr 8, 2024 |  |  |  |  |  |
| StLucia-GrosPiton | *T. mesoleucus* | Apr 16 - Apr 20, 2015 | 8 | 32 | 13.81 | -61.06 | 293 |
| StLucia-VillaPiton |  | Mar 31 - Apr 4, 2024 |  |  |  |  |  |
| StVincent-Buccament Valley | *T. musicus* | Apr 11 - Apr 15, 2015 | 6 | 28 | 13.19 | -61.24 | 206 |
| StVincent-MtVernon |  | Mar 25 - Mar 30, 2024 |  |  |  |  |  |
| Grenada-StGeorge's | *T. grenadensis* | Apr 6 - Apr 10, 2015 | 9 | 30 | 12.10 | -61.74 | 256 |
| Grenada-StDavids |  | Mar 21 - Mar 24, 2024 |  |  |  |  |  |
| Tobago-Castara | *T. m. tobagensis* | Apr 1 - Apr 5, 2015 | 8 | 30 | 11.28 | -60.69 | 95 |
| Tobago-Castara |  | Mar 17 - Mar 20, 2024 |  |  |  |  |  |
| Trinidad-Maravel Valley | *T. m. clarus* | Apr 27 - Apr 30, 2015 | 10 | 31 | 10.69 | -61.29 | 245 |
| Trinidad-Arima Valley |  | Mar 13 - Mar 16, 2024 |  |  |  |  |  |
| Ecuador-SantoDomingo | *T. m. albicans* | Apr 7 - Apr 8, 2022 | 2 | 10 | 0.27 | -79.14 | 611 |
| Ecuador-Maquipacuna | *T. m. albicans* | Mar 23 - Mar 25, 2022 | 3 | 8 | 0.13 | -78.63 | 1,367 |
| Ecuador-Mindo | *T. m. albicans* | Mar 26 - Mar 28, 2022 | 5 | 22 | 0.03 | -78.80 | 1,141 |
|  |  | Apr 8 - Apr 10, 2022 |  |  |  |  |  |
| Ecuador-Milpes | *T. m. albicans* | Mar 29 - Mar 30, 2022 | 2 | 8 | 0.03 | -78.87 | 1,150 |
| Ecuador-CopaLinga | *T. m. clarus* | Apr 1 - Apr 4, 2022 | 7 | 16 | -4.09 | -78.96 | 1,017 |

Specifically, Klicka et al. (2023) hypothesized that birds on Dominica might have originated from Central America while those on the other islands had a south American origin. It is possible that the patterns of similarity and difference in songs among the islands, and between the various islands and the mainland populations, can furnish additional evidence with which to evaluate this hypothesis. Table 1 provides further details on the sample populations and their specific locations.

### Song Recording

House wrens are noted for their variable (complex) song patterns (Kroodsma 1977; Tubaro 1990; Rendall and Kaluthota 2013; dos Santos et al 2016). In this study, every effort was therefore made to capture and represent that variability but also to standardize recordings to eliminate as many sources of extraneous or confounding influence as possible.

To that end, all research and recording was done by the same individual (DR), using the same equipment and recording parameters: a Sound Devices 702 (or 722) digital recorder and Telinga Twin Science Pro6/8/9-MK2 microphones, with corresponding 24” parabola, using a digital sampling rate of 48kHz and 24 bit depth. Research was also conducted at the same time of year for all populations (March – April), in spring, to control as best possible for seasonal and breeding stage variation in song patterns. This is harder to ensure for resident, tropical populations compared to migratory populations in the temperate zone, because tropical populations are often not as seasonally synchronized in their breeding activity. However, previous research on tropical House Wrens in Costa Rica (Young 1994) shows that for this species there is nevertheless a breeding peak in early spring (March – May: Young 1994) similar to that in the north temperate zone.

I also focused on singing only during the dawn chorus to further control for variation related to breeding stage but also to minimize diel variation in singing activity. Dawn is when singing is most vigorous and sustained for many passerines, and singing often decreases considerably thereafter.

Previous focused research on the northern House Wren confirms this and shows the early morning hours to be the time of day when male singing is at its peak of both performance and complexity and also when the differences between the song patterns of paired and unpaired males are minimal (Kaluthota et al. 2020).

Recordings at every location focused on obtaining a sample that balanced sampling different males in the population but also capturing variation in singing activity within individual males which is known for continental populations of house wrens to be substantial and biologically relevant (Johnson and Kermott 1991; Johnson and Searcy 1996; Deslandes et al. 2014; dos Santos et al. 2018; DiSciullo et al. 2024).

### Song Sample

The song sample selected for detailed analysis in this study was compiled from the larger database of recordings obtained from 92 days of field observation across 20 different sample locations. Songs were selected from this larger database using a number of complementary criteria. First, songs had to be of the highest possible quality lacking background noise and overlapping song or vocalizations from other species. The focus of further selection was then on representing population level variation as best possible by including songs from multiple different males from each location but also multiple songs from individual males. The latter dimension is important because previous work on continental populations of both northern and southern House Wren shows that individual males produce a large repertoire of notes and syllables that they combine into even larger repertoires of different song types but that their often repetitive singing style means that much of this variation is hidden in the short term (Kroodsma 1977; Tubaro 1990; Rendall and Kaluthota 2013; dos Santos et al 2016); hence it it takes some time to capture the variation of which individual males are capable. To reflect both dimensions of variation, the sample for each location attempted to balance the representation of different males with contributions from individual males. At the same time, to avoid over-representing any specific males, no male contributed more than five songs to the sample for any location. The final and important selection criterion to properly represent population level song variation was that every song included in the sample, whether from the same male or a different male, had to be of a different song type, as commonly defined in song research and as implemented in previous detailed characterizations of song for both northern and southern House Wren (Rendall and Kaluthota 2013; dos Santos et al. 2016) as the specific sequence of different notes and syllables. There was no point including multiple versions of the same song type; this would simply involve a form of pseudoreplication that might bias the sample for some locations by under-representing the range of variation. Hence, every song in the sample represents a different song type – a different sequence of notes and syllables. The sample ultimately compiled for detailed analysis involved 293 songs from 85 different males. Table 1 provides a more detailed breakdown of the sample by location.

### Song Analysis

Preview of song recordings indicated that songs of house wrens in the Lesser Antilles follow the same basic organizational structure and patterns of delivery that has has been reported in previous detailed studies of other house wren populations (Kroodsma 1977; Tubaro 1990; Rendall and Kaluthota 2013; Sosa-López and Mennill 2014a, b, c; dos Santos et al. 2016). Hence, to facilitate comparisons between studies and among all populations, the analysis of songs studied here followed the same general approach and methods used previously, and involved two different stages.

The first stage involved qualitative characterization of the relatively high-level patterns of song structure, organization and delivery, including how songs are constructed from constituent notes and syllables (defined as regularly occurring combinations of 2 or more notes); how they are organized into different song types (defined as a different sequence of the individual notes and syllables); how singing is patterned into bouts; and how frequently successive songs in a bout involve repeating or changing song types.

The second stage focused on the more detailed structure of individual songs and involved quantification of 21 different temporal and spectral features encompassing both the Introduction and Main sections of songs (see Table 2 and Figure 2). Measured song parameters were: the overall duration of each song (SONG-Duration) and the duration of its Introduction (INTRO-Duration) and Main sections (MAIN-Duration); the number of elements in the entire song (SONG-#Elements) and in the Introduction and Main sections (INTRO-#Elements, MAIN-#Elements); the average duration of the elements and the intervals between them in each section (INTRO-ElementDuration; INTRO-IntervalDuration; MAIN-ElementDuration; MAIN-IntervalDuration); the rate of element production across the entire song (ElemProdRate); the center frequency of elements in the Introduction and Main sections (INTRO-CenterFreq, MAIN-CenterFreq); the average signal-to-noise ratio of elements in the Introduction (INTRO-SNR); the average entropy of elements in the Introduction and Main sections (INTRO-AvgEntropy, MAIN-AvgEntropy); the average bandwidth of elements in the Main section (MAIN-BW-Element); the mean fundamental frequency (MeanF_0_) and extent of frequency modulation of each element in the Main section (MeanFM/s); the overall duty cycle of each song, defined as the proportion of each song allocated to active song (sound production) versus silent gaps (SONG-DutyCycle); and the proportion of each song represented by the Main section (Main-%Duration).

**Figure 2:**
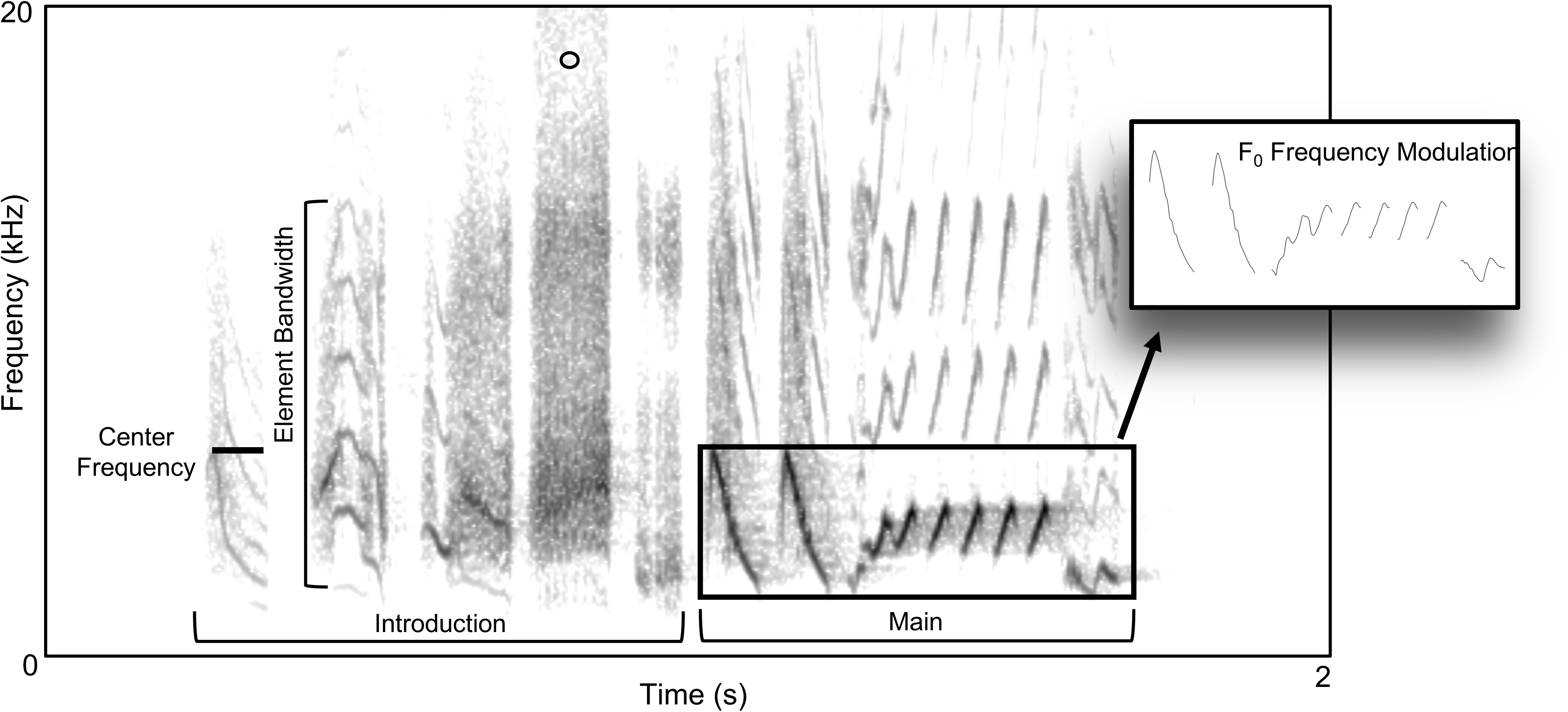
Spectrogram of standard house wren song variant. This example is from Grenada and illustrates the standard structural organization of song common to all populations involving two distinct sections: an Introductory section of primarily noisy broad-band notes, or more tonal notes with a noisy overlay; and a Main section of more distinctive, tonal and frequency modulated notes and syllables. Many, but not all, songs include a conspicuous buzzy note (o) at or near the transition between the two sections. Some of the other measured features of song are also labelled.

**Table 2.**
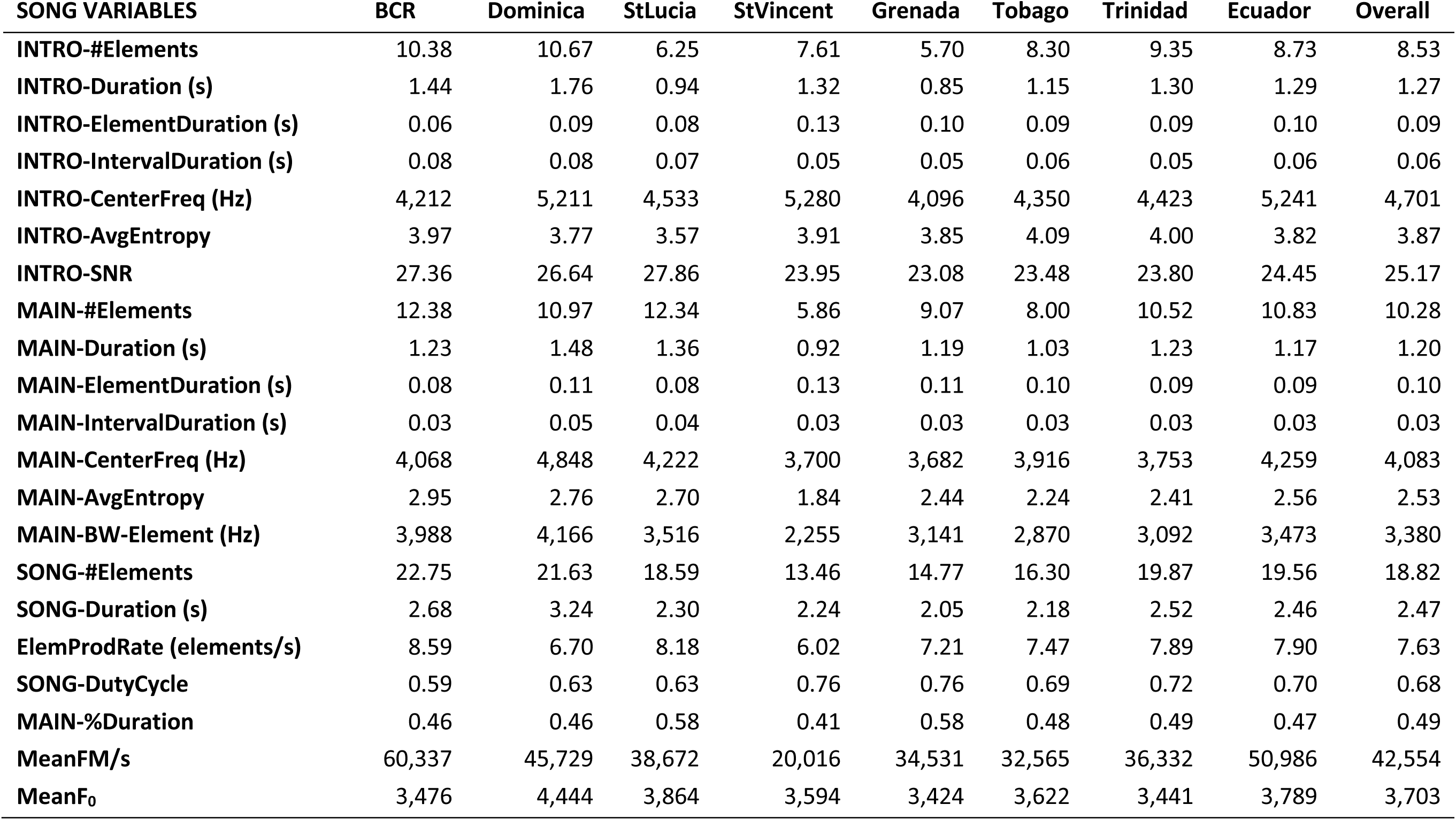
Means for song variables by population.

| SONG VARIABLES | BCR | Dominica | StLucia | StVincent | Grenada | Tobago | Trinidad | Ecuador | Overall |
| --- | --- | --- | --- | --- | --- | --- | --- | --- | --- |
| INTRO-#Elements | 10.38 | 10.67 | 6.25 | 7.61 | 5.70 | 8.30 | 9.35 | 8.73 | 8.53 |
| INTRO-Duration (s) | 1.44 | 1.76 | 0.94 | 1.32 | 0.85 | 1.15 | 1.30 | 1.29 | 1.27 |
| INTRO-ElementDuration (s) | 0.06 | 0.09 | 0.08 | 0.13 | 0.10 | 0.09 | 0.09 | 0.10 | 0.09 |
| INTRO-IntervalDuration (s) | 0.08 | 0.08 | 0.07 | 0.05 | 0.05 | 0.06 | 0.05 | 0.06 | 0.06 |
| INTRO-CenterFreq (Hz) | 4,212 | 5,211 | 4,533 | 5,280 | 4,096 | 4,350 | 4,423 | 5,241 | 4,701 |
| INTRO-AvgEntropy | 3.97 | 3.77 | 3.57 | 3.91 | 3.85 | 4.09 | 4.00 | 3.82 | 3.87 |
| INTRO-SNR | 27.36 | 26.64 | 27.86 | 23.95 | 23.08 | 23.48 | 23.80 | 24.45 | 25.17 |
| MAIN-#Elements | 12.38 | 10.97 | 12.34 | 5.86 | 9.07 | 8.00 | 10.52 | 10.83 | 10.28 |
| MAIN-Duration (s) | 1.23 | 1.48 | 1.36 | 0.92 | 1.19 | 1.03 | 1.23 | 1.17 | 1.20 |
| MAIN-ElementDuration (s) | 0.08 | 0.11 | 0.08 | 0.13 | 0.11 | 0.10 | 0.09 | 0.09 | 0.10 |
| MAIN-IntervalDuration (s) | 0.03 | 0.05 | 0.04 | 0.03 | 0.03 | 0.03 | 0.03 | 0.03 | 0.03 |
| MAIN-CenterFreq (Hz) | 4,068 | 4,848 | 4,222 | 3,700 | 3,682 | 3,916 | 3,753 | 4,259 | 4,083 |
| MAIN-AvgEntropy | 2.95 | 2.76 | 2.70 | 1.84 | 2.44 | 2.24 | 2.41 | 2.56 | 2.53 |
| MAIN-BW-Element (Hz) | 3,988 | 4,166 | 3,516 | 2,255 | 3,141 | 2,870 | 3,092 | 3,473 | 3,380 |
| SONG-#Elements | 22.75 | 21.63 | 18.59 | 13.46 | 14.77 | 16.30 | 19.87 | 19.56 | 18.82 |
| SONG-Duration (s) | 2.68 | 3.24 | 2.30 | 2.24 | 2.05 | 2.18 | 2.52 | 2.46 | 2.47 |
| ElemProdRate (elements/s) | 8.59 | 6.70 | 8.18 | 6.02 | 7.21 | 7.47 | 7.89 | 7.90 | 7.63 |
| SONG-DutyCycle | 0.59 | 0.63 | 0.63 | 0.76 | 0.76 | 0.69 | 0.72 | 0.70 | 0.68 |
| MAIN-%Duration | 0.46 | 0.46 | 0.58 | 0.41 | 0.58 | 0.48 | 0.49 | 0.47 | 0.49 |
| MeanFM/s | 60,337 | 45,729 | 38,672 | 20,016 | 34,531 | 32,565 | 36,332 | 50,986 | 42,554 |
| MeanF<sub>0</sub> | 3,476 | 4,444 | 3,864 | 3,594 | 3,424 | 3,622 | 3,441 | 3,789 | 3,703 |

Measurements were made using both Raven Pro (V1.6.1, Cornell Lab of Ornithology 2019) and PRAAT (V6.1.4, Boersma & Weenink 2021) software using the following spectrogram parameters: Hann Window, with 3db (135 Hz) filter bandwidth, 512 point DFT, and 90.2% overlap in successive spectral frames (or a 50 point, or 1.04 ms, “hop” size).

### Statistical Analysis

The first phase of song analysis was qualitative in nature and therefore did not involve application of statistical procedures. For the data in the second, quantitative phase, multivariate discriminant analysis was used to test the extent to which the detailed features of songs were discriminably different among the various islands of the Lesser Antilles, as well as from mainland forms, in alignment with the recent taxonomic reclassification of the island populations as distinct species. Discriminant analysis is notably sensitive to the number of predictor variables used to discriminate among groups, where a high ratio of predictor variables to groups can produce unstable solutions that ‘over-fit’ the data and thereby inflate discriminability (Klecka 1980, Tabachnick & Fiddell 2007). To avoid this problem, a Principle Components Analysis (PCA) was first conducted on the complete set of 21 measured acoustic features, with no rotation of the factor solution. This precautionary step allowed examination of natural covariation among the original acoustic features according to their associations within and between PCA factors. It also importantly reduced the number of predictor variables to be used in discriminant analysis to a much smaller set of orthogonal multivariate dimensions that nevertheless retained most of the variation in the entire set of original acoustic features. The practical success of the discriminant solution was evaluated using a jacknife, leave-one-out classification procedure which proceeds serially to classify each case in turn based on functions derived from all of the other cases except the one that is currently being classified (the one left out). It is a fairly liberal cross-validation technique but avoids the circularity of including in the derivation of the classification functions the very cases that are then to be classified from those functions. PCA and discriminant analysis were implemented in both SPSS (version 31.0) and NCSS (2023).

## RESULTS

### General Patterns of Song Structure, Organization and Delivery

In general patterns of structure, organization and delivery, male song is similar across all sampled populations and follows closely patterns described previously for continental forms of northern and southern House Wren. Thus, it is a fast-paced jumble of variably structured notes that is nevertheless predictably organized in two distinct sections: an Introductory section of broadband notes that can be harsh and unstructured, or more tonal with harmonic overtones or a noisy overlay, many resembling or replicating common call notes produced in other contexts; and a terminal or Main section comprising much more distinctive and clearly structured tonal and frequency-modulated notes and syllables. Introductory elements are often repeated multiple times, creating a ’stuttering’ effect, and there is often, but not always, a distinctive and emphatic ‘buzzy’ note at the junction between the Introduction and Main sections of each song (see Figure 2). The terminal or Main section of the song is typically considerably louder than the Introduction and can be heard over much longer distances.

While most songs conform to this standard Introduction-Main template, males in all populations also show a capacity to deviate from it. Thus, songs occasionally omit one or other section, or involve one or more introductory-type notes appended to the end of the Main section of a song, or an entire second Main section appended following a very short gap only marginally longer than the typical interval between syllables within a song (what might be labelled a double-Main song). Songs can also entail a protracted concatentation of two or three complete Introductory-Main sections with virtually no gap demarcating individual songs, what might be labelled song ‘doublets’ or ‘triplets’. The capacity for such variety is present in all populations, but its frequency of use appears to vary among some of them.

Males in all populations also have a large repertoire of Introductory and Main notes and syllables that they recombine to produce a much larger repertoire of different song types. In bouts of singing, males typically repeat the same song type many times before switching to a different type, although they commonly vary the number of repetitions of particular syllabe types within each song from one rendition to the next. And when they switch song types, they tend to do so incrementally by the addition, deletion, substitution, or movement of only one or two syllable types at a time. The common pattern of song delivery then is one of ‘eventual variety’ where the diversity of a male’s repertoire of different song types is displayed only gradually (sensu Kroodsma 1977). However, males in all populations also show a capacity for more wholesale changes in successive song types and for more rapid switching amomg them, thereby sometimes manifesting a pattern of ‘immediate variety’ in their singing (see Figure 3 for example; see also Figure 1 in Juárez et al. 2025). The latter delivery style is more common when males are close to and courting a female, or responding to song from other males.

**Figure 3:**
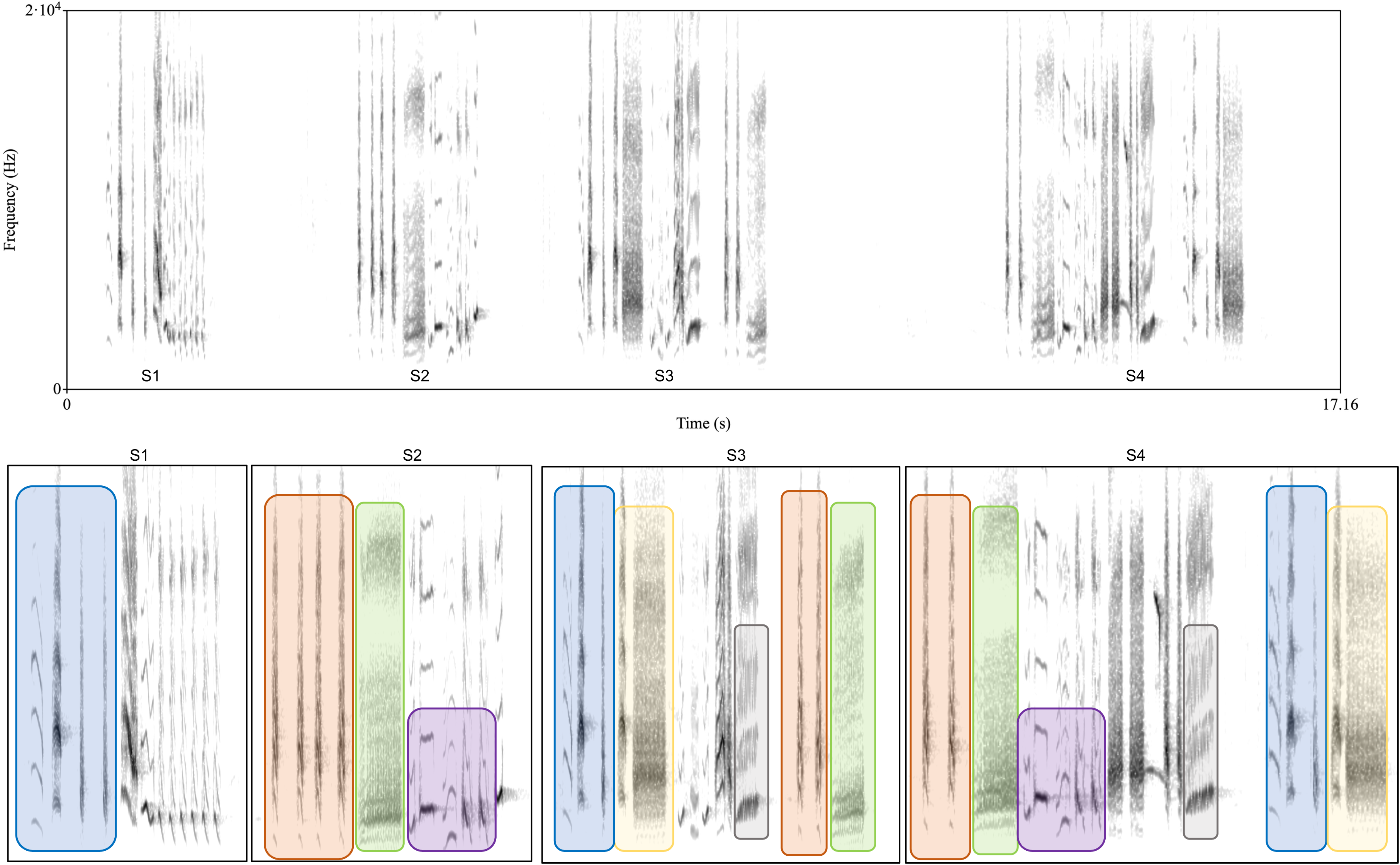
A short sequence of four songs from a longer bout of singing from St Lucia. This short sequence illustrates several notable features of song organization and song type variation. The differently sized and colored boxed segments are used to highlight a specific note or note combination (syllable) to help illustrate the patterns. The typical pattern of song organization and delivery in St Lucia and elsehere commonly entails repeating the same song type many times before switching (eventual variety) and then changing song types only gradually by the addition, deletion, substitution, or movement of one or two notes or syllables at a time. However, this example illustrates the capacity for more immediate variety in, and some wholesale changes between, different song types that can occur in all populations but may be elaborated to a greater degree in St Lucia. Thus, S1 and S2 are completely different in both their Introduction and Main sections. S3 uses the first three notes from the Introduction of S1 (blue box) and adds a buzzy element (yellow box) to the Introduction and then introduces another different Main section with a rising buzz note (grey box) at the end of the Main section; it then also appends Introductory notes from S2 (orange and green boxes) to the end of the first canonical song after a very short gap. S4 reverts to the Introductory notes and most of the Main section from S2, adds some new buzz and short tick notes and the rising buzz note from S3 (grey box), and then appends to the end of the song, after a long gap, the Introduction from S3 (blue and yellow boxes), the first part of which (blue box) was also the Introduction to S1. In addition to thereby illustrating more immediate variety in successive song types, this example also illustrates how songs (particularly in St Lucia) sometimes involve greater continuous mixing of Introduction and Main type notes (S4) or simply appending Introduction type notes to the end of the Main section of a song after a short gap (as in S3 and S4).

The diel pattern of singing is also similar across all sampled populations and follows a pattern typical of tropical species generally. It involves a relatively short period of vigorous singing at dawn, lasting 10-30 minutes, sometimes preceded by a few tentative songs (of lower amplitude, and relatively short duration with longer intervals between songs) in the hour pre-dawn. Bouts of singing at dawn typically involve high rates of song production (8-12 songs/minute). For the remainder of the day, males produce short bouts of 2-4 songs sporadically as they move about their territory in the course of routine activities. The exception is that males can sing far more vigorously and continously throughout the day when actively courting a female.

While broadly similar in the above-noted general patterns of song organization and delivery, there is nevertheless some variation in their manifestation among populations. Thus songs of males in St Vincent appear to adhere much more strictly to the basic Introduction-Main template and manifest fewer deviations from it, and appear aslo to involve an exaggerated level of song type repetition: for example, one unpaired male was observed (and recorded) to repeat the same song type 102 times in succession over a 13-min interval, and to produce only four slightly different song types over the course of 214 successive songs; on another occasion, he repeated the same song type 79 times before switching. In contrast, the organization of song by males in St Lucia appears to be more flexible with less delineation between, and more fluid mixing among, Introductory and Main type notes and syllables. Males in St Lucia also sometimes sing protracted and highly variable sequences of Introductory and Main type notes and syllables with no detectable gaps demarcating individual songs in a pattern of continuous delivery reminiscient of some members of the *Mimidae*, such as the north American Catbird (*Dumetellus*).

There is also variation in the form and diversity of notes and syllables among some of the populations. Male song in St Vincent is notably distinctive in this respect: introductory elements include noisy broadband notes as elsewhere but a greater prevelance of higher-frequency notes that often entail two distinct tones that are harmonically unrelated to each other, which gives the introductory section of many songs a uniquely shrill and raspy, or even discordant, quality. Notes in the Main section of the song in St Vincent are also notably distinct in being limited in the number and variety of forms they take (see example in Figure 4). These include a few virtually pure-tone whistles, which are not common in other populations, and a limited variety of only subtly different narrow-band, chevron and inverted chevron-shaped notes. The latter are typically produced in pairs with modest frequency shifts, or offsets, between notes in a pair (e.g., **^ v; ^** _^_ ; _v_ **^v^** ) which creates a highly distinctive alternating high-low, sing-song quality ("*see-soo, see-soo*") that may motivate the latin name for the species (*musicus*). At the same time, there are few broad-band frequency sweeps or protracted trills in the Main section of song in St Vincent which are relatively common features of song in the other populations. The latter broad-band frequency sweeps are especially common and elaborated in the Main section of songs in Dominica which tend to be longer and to involve a preponderance of such sweeps, many spanning 5 kHz or more, organized in extended trills (see example in Figure 5).

**Figure 4:**
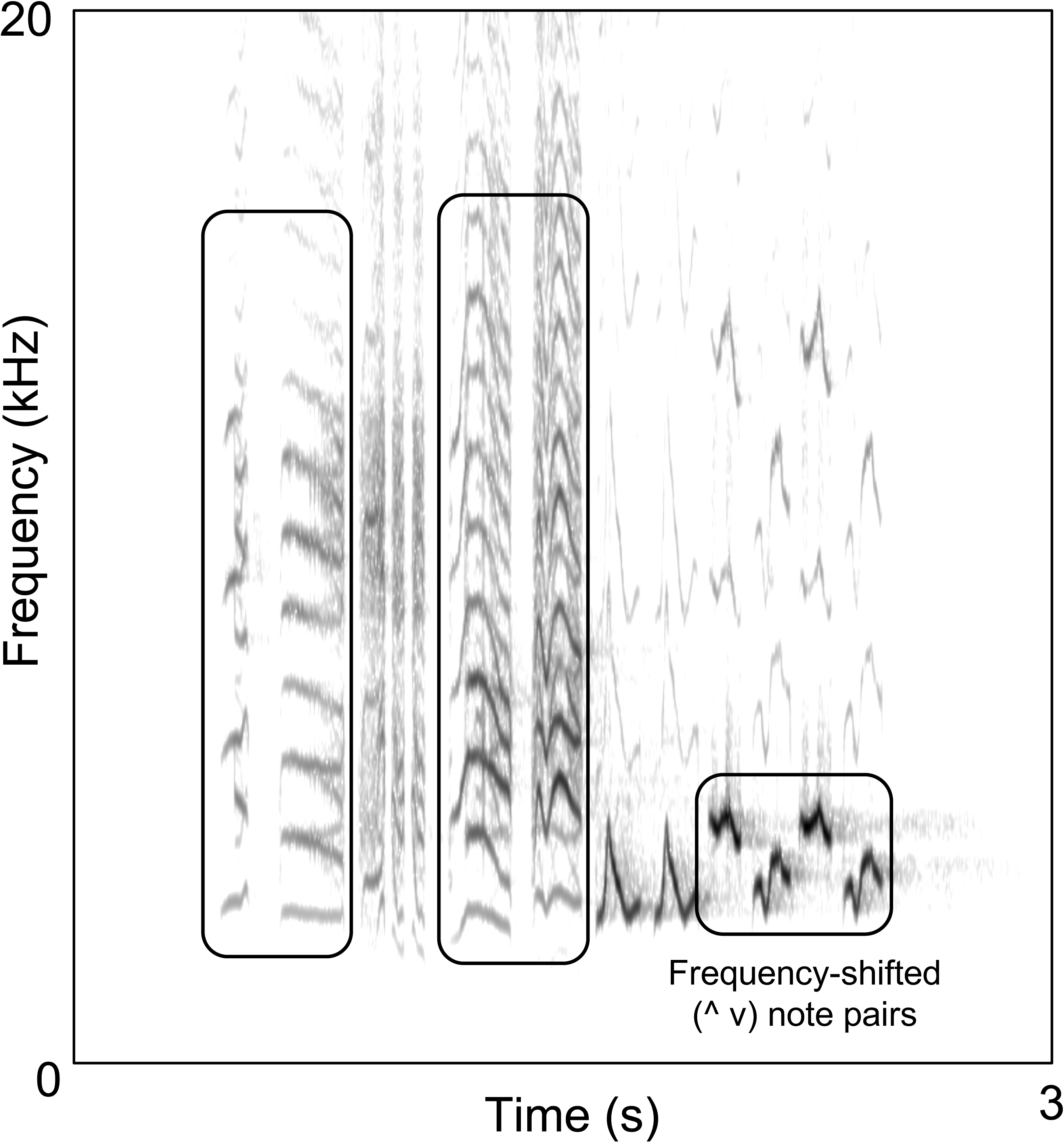
A representative song from St Vincent illustrating some of the distinctive features of song from this island population. These include higher frequency elements in the Introduction section with harmonically unrelated tonal components (tall rectangular boxes) and a comparatively short and simple Main section comprised of narrow band, chevron shaped notes (^v) often produced in pairs and offset slightly in frequency producing a distinctive high-low (‘sing-song’) percept.

**Figure 5:**
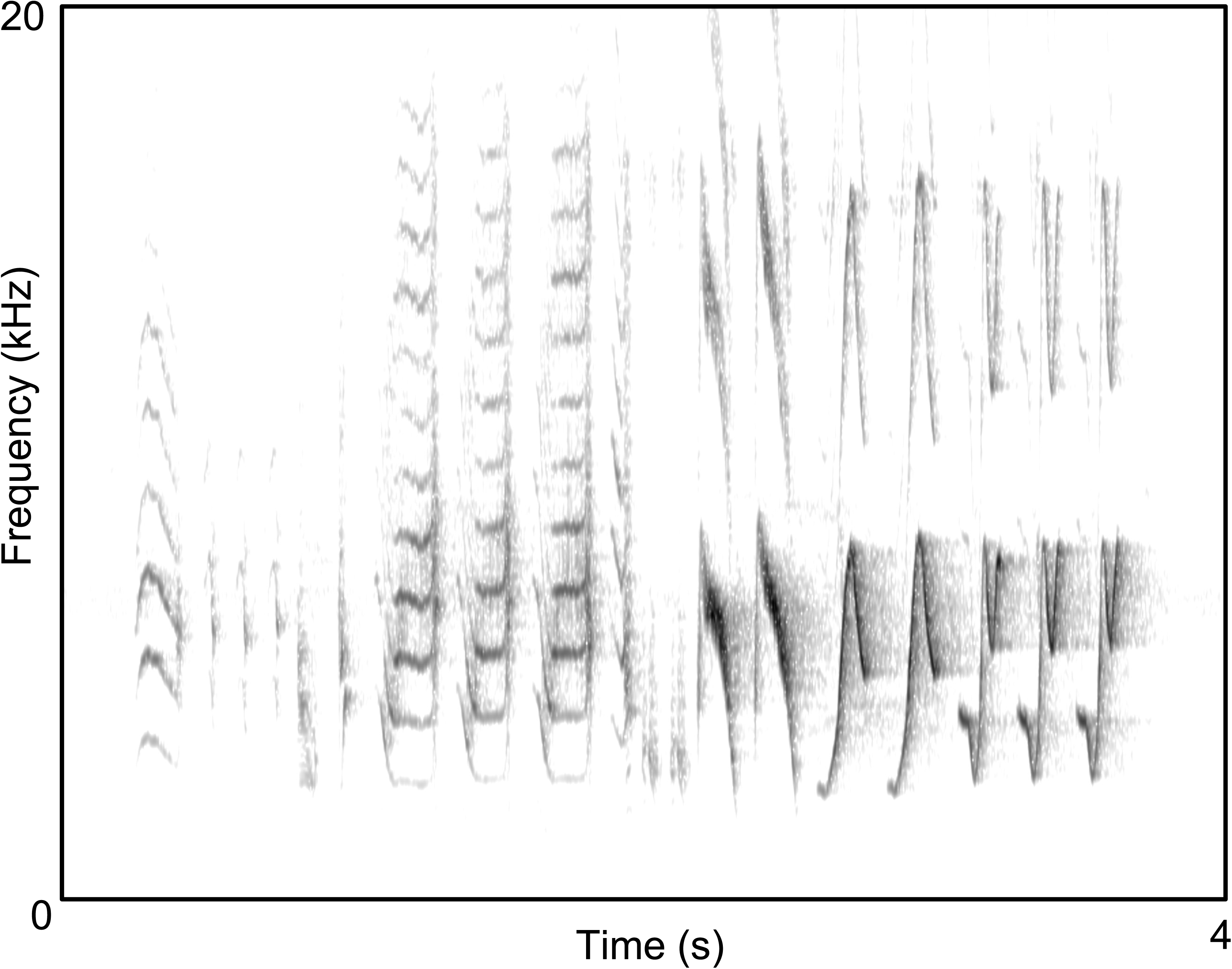
A representative song from Dominica illustrating the more widely spaced elements and preponderance of very wide-band frequency sweeps in the Main section.

### Detailed Song Structure

Mean values for each of the 21 detailed song features are provided for each population in Table 2. Inspection of these values confirms some of the differences among populations just noted: for example, that songs in St Vincent have notably shorter, simpler Main sections, comprised of very few elements that also have the lowest levels of frequency modulation (FM/s); while songs in Dominica are notably longer than all of the other populations, with the longest Main section comprised of elements with comparatively high levels of frequency modulation.

Principle Components Analysis on this set of 21 acoustic features produced a smaller set of seven orthogonal factors, or components, with eigenvalues greater than one that cumulatively accounted for and retained 83.9% of the variation in the complete dataset (Table 3). Fully two-thirds of this variation was accounted for by the first three components. The association of original variables with the different components is detailed in Table 4.

**Table 3.** Summary of principle components analysis.

| Component | Eigenvalue | % of Variance | Cumulative % |
| --- | --- | --- | --- |
| **1** | 5.584 | 26.591 | 26.591 |
| **2** | 3.465 | 16.501 | 43.092 |
| **3** | 2.693 | 12.823 | 55.914 |
| **4** | 1.958 | 9.324 | 65.238 |
| **5** | 1.458 | 6.943 | 72.182 |
| **6** | 1.280 | 6.096 | 78.277 |
| **7** | 1.186 | 5.650 | 83.927 |

**Table 4.** Variable loadings on each component from principle components analysis.

| Acoustic variables | Component* |  |  |  |  |  |  |
| --- | --- | --- | --- | --- | --- | --- | --- |
|  | 1 | 2 | 3 | 4 | 5 | 6 | 7 |
| INTRO-#Elements | 0.436 | **0.609** | -0.555 | 0.166 | -0.011 | 0.074 | -0.246 |
| INTRO-Duration | 0.318 | **0.770** | -0.427 | 0.283 | 0.015 | 0.014 | 0.106 |
| INTRO-ElementDuration | **-0.647** | 0.142 | 0.135 | 0.141 | -0.097 | 0.263 | 0.394 |
| INTRO-IntervalDuration | 0.417 | 0.031 | 0.309 | -0.006 | 0.176 | -0.571 | **0.509** |
| INTRO-CenterFreq | -0.157 | 0.292 | 0.197 | -0.002 | -0.207 | 0.276 | **0.570** |
| INTRO-AvgEntropy | 0.013 | -0.070 | -0.131 | 0.080 | **0.772** | 0.085 | 0.202 |
| INTRO-SNR | 0.212 | 0.263 | 0.116 | -0.286 | **-0.624** | -0.337 | -0.055 |
| MAIN-#Elements | **0.737** | -0.523 | 0.125 | 0.280 | -0.147 | 0.137 | 0.149 |
| MAIN-Duration | 0.489 | -0.325 | 0.512 | **0.597** | -0.045 | 0.095 | -0.074 |
| MAIN-ElementDuration | **-0.637** | 0.425 | 0.288 | 0.228 | 0.240 | -0.069 | -0.264 |
| MAIN-IntervalDuration | -0.112 | 0.466 | **0.566** | 0.299 | -0.023 | -0.270 | -0.145 |
| MAIN-CenterFreq | 0.342 | 0.457 | **0.505** | -0.336 | -0.048 | 0.342 | -0.071 |
| MAIN-AvgEntropy | **0.744** | 0.050 | 0.220 | -0.273 | 0.306 | 0.080 | -0.034 |
| MAIN-BW-Element | 0.365 | 0.357 | **0.526** | -0.255 | 0.250 | 0.043 | -0.349 |
| SONG-#Elements | **0.876** | -0.039 | -0.250 | 0.333 | -0.127 | 0.158 | -0.035 |
| SONG-Duration | 0.609 | 0.402 | 0.013 | **0.657** | -0.019 | 0.079 | 0.035 |
| ElemProdRate | **0.620** | -0.509 | -0.364 | -0.286 | -0.153 | 0.118 | -0.116 |
| SONG-DutyCycle | **-0.756** | -0.160 | -0.034 | 0.135 | -0.018 | 0.505 | -0.122 |
| MAIN-%Duration | 0.082 | **-0.739** | 0.605 | 0.174 | -0.016 | 0.043 | -0.130 |
| MeanFM/s | **0.695** | 0.082 | -0.007 | -0.429 | 0.257 | 0.176 | 0.162 |
| MeanF<sub>0</sub> | 0.199 | **0.450** | 0.442 | -0.238 | -0.210 | 0.343 | 0.107 |

\*Bolted values indicate for each acoustic variable the component with which they are most strongly associated.

Results of Discriminant Analysis using components from PCA are provided in Table 5. The analysis produced seven discriminant functions (one for each component from PCA), all but the last of which contributed some degree of discrimination among the populations. However, the bulk of the discrimination was carried by the first two functions (DF1 and DF2) which each had eigenvalues > 1.0, markedly higher Canonical Correlations and lower Wilk’s Lambda scores than the other functions, and together accounted for 84.3% of the variation in all of the components from PCA. DF3 and DF4 provided notably less, but not trivial, levels of discrimination. The associations of individual components from PCA with each discriminant function are shown in the structure matrix in Table 6.

**Table 5.** Summary results of discriminant analysis on principle component scores.

| Function | Eigenvalue | %<br>Variance | Cumulative<br>% Variance | Canonical<br>Correlation | Wilk's<br>Lambda | F-Value | P-value |
| --- | --- | --- | --- | --- | --- | --- | --- |
| DF1 | 3.346 | 63.1 | 63.1 | 0.877 | 0.052 | 22.9 | <0.001 |
| DF2 | 1.123 | 21.2 | 84.3 | 0.727 | 0.226 | 13.8 | <0.001 |
| DF3 | 0.385 | 7.3 | 91.5 | 0.527 | 0.480 | 9.1 | <0.001 |
| DF4 | 0.266 | 5.0 | 96.5 | 0.458 | 0.664 | 7.7 | <0.001 |
| DF5 | 0.143 | 2.7 | 99.2 | 0.353 | 0.841 | 5.6 | <0.001 |
| DF6 | 0.039 | 0.7 | 100.0 | 0.194 | 0.961 | 2.9 | 0.0226 |
| DF7 | 0.002 | 0.0 | 100.0 | 0.041 | 0.998 | 0.5 | 0.4871 |

**Table 6.**
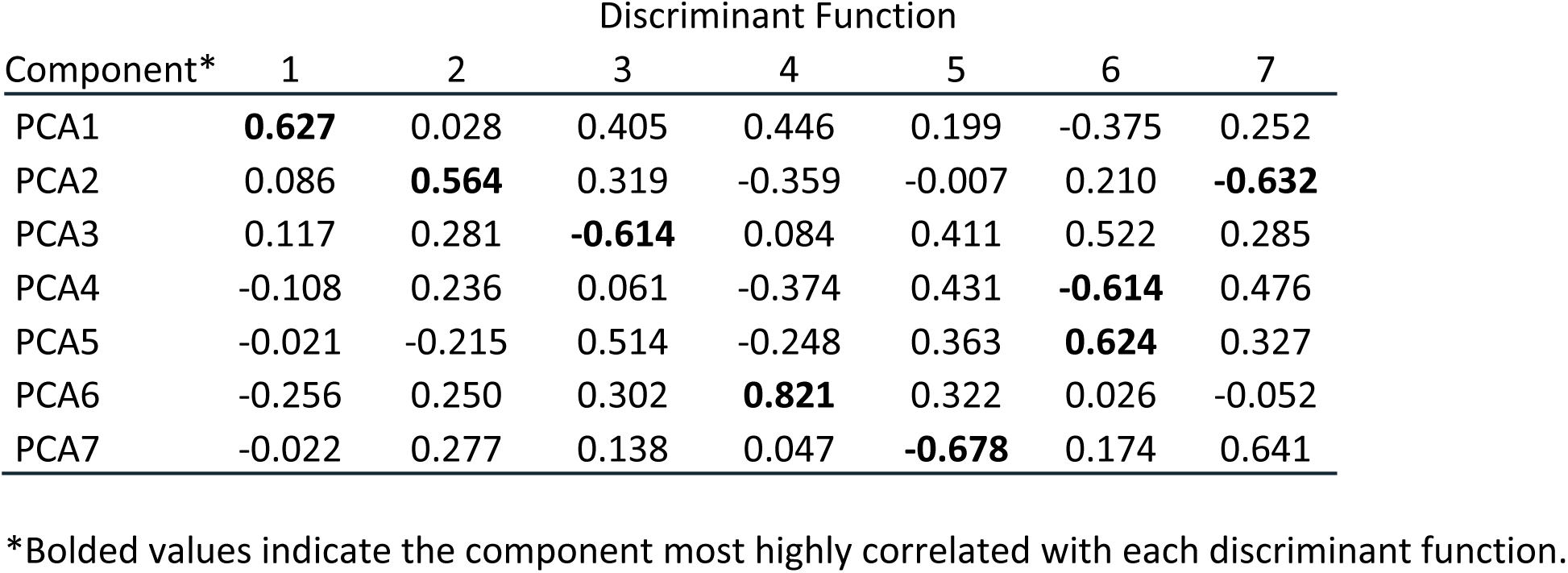
Structure matrix from discriminant analysis showing the correlations of individual principle components with each discriminant function.

Discriminant scores for each song are shown in bivariate plots for the first three discriminant functions in Figure 6. The first panel, plotting DF1 against DF2, encompasses almost 85% of the variation in the data and shows some separation of BCR, Dominica, and St Vincent and to a much lesser extent Grenada, with the remaining populations clustered centrally with highly overlapping or virtually coincident distributions: Ecuador at the center and Trinidad and Tobago immediately adjacent to that and co-located. The second panel, plotting DF1 against DF3, shows that DF3 helps to separate St Lucia from the central nexus. It also highlights a higher-level pattern of population clustering that groups BCR, Dominica and St Lucia together and separately from a clustering of the remaining populations. The third panel (DF2 vs DF3) shows that there is little further separation of populations gained from additional functions.

**Figure 6.**
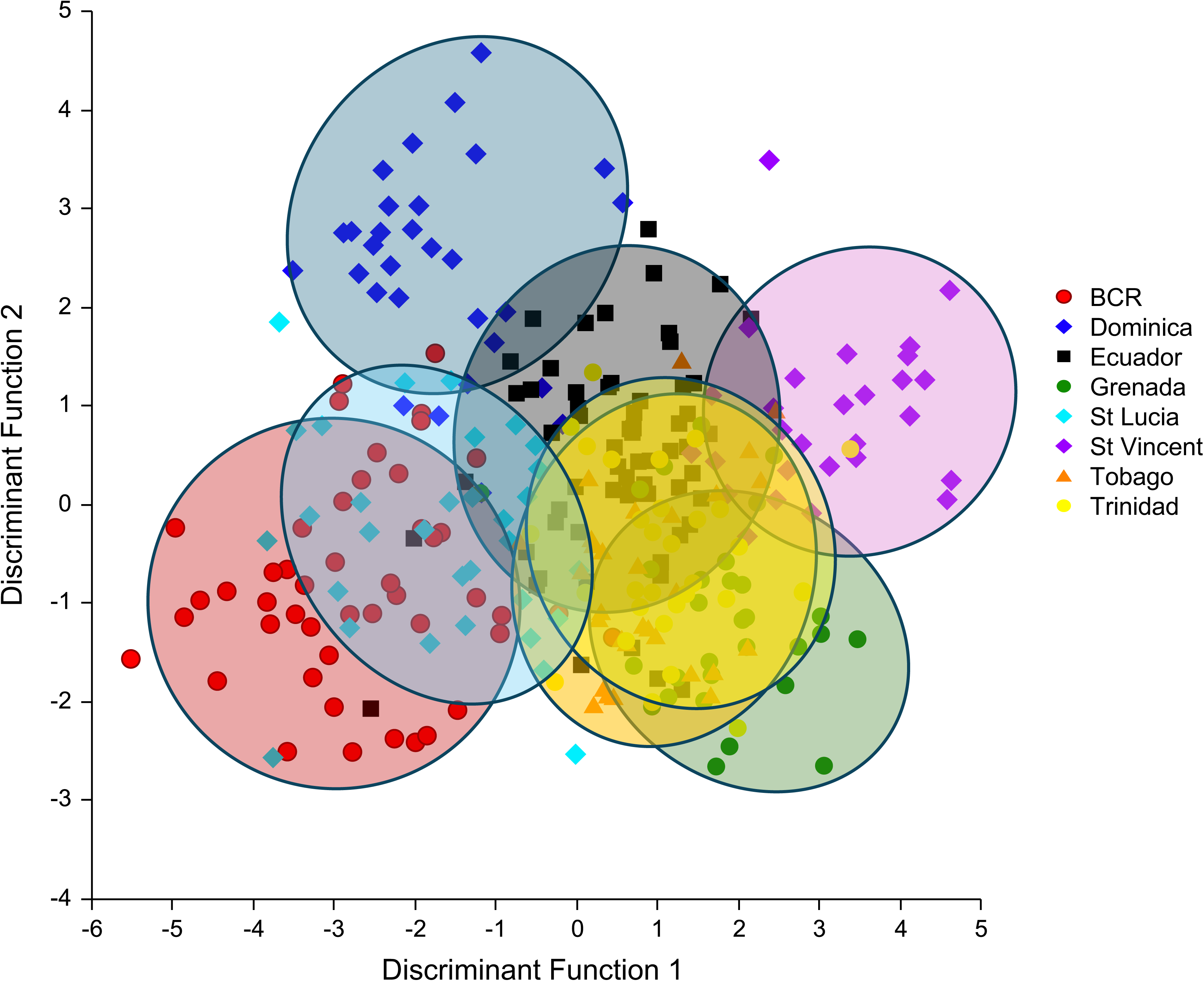
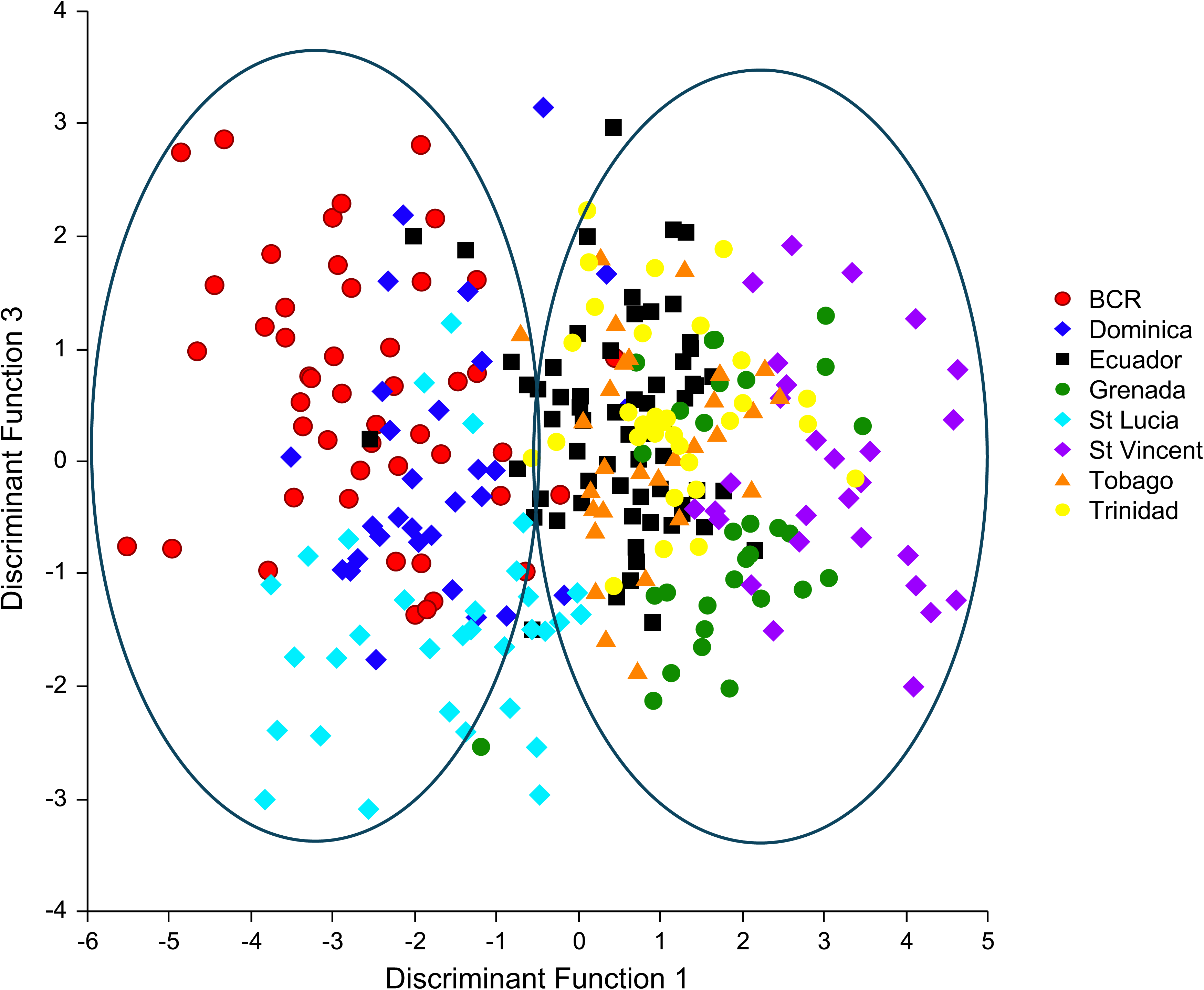
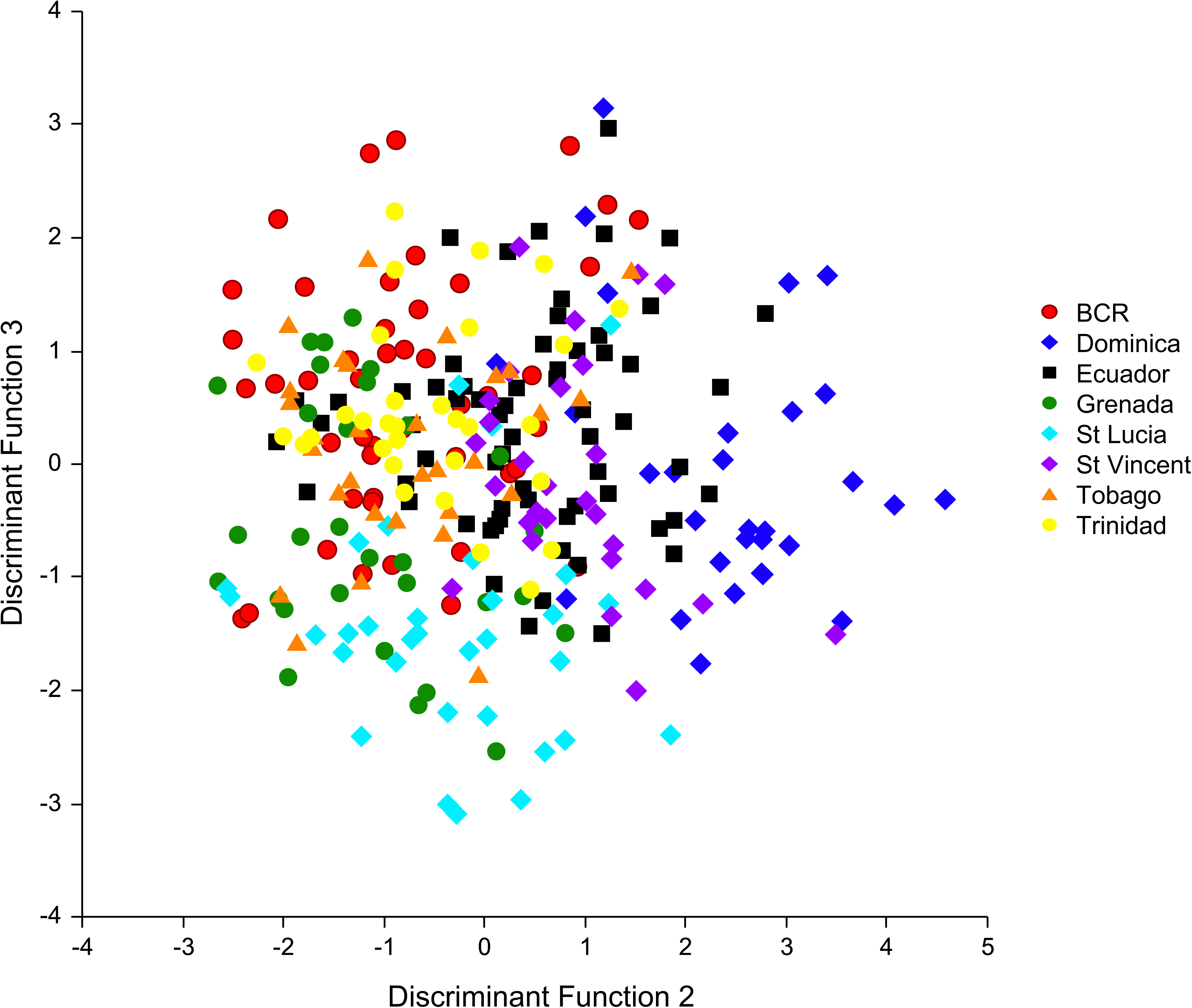
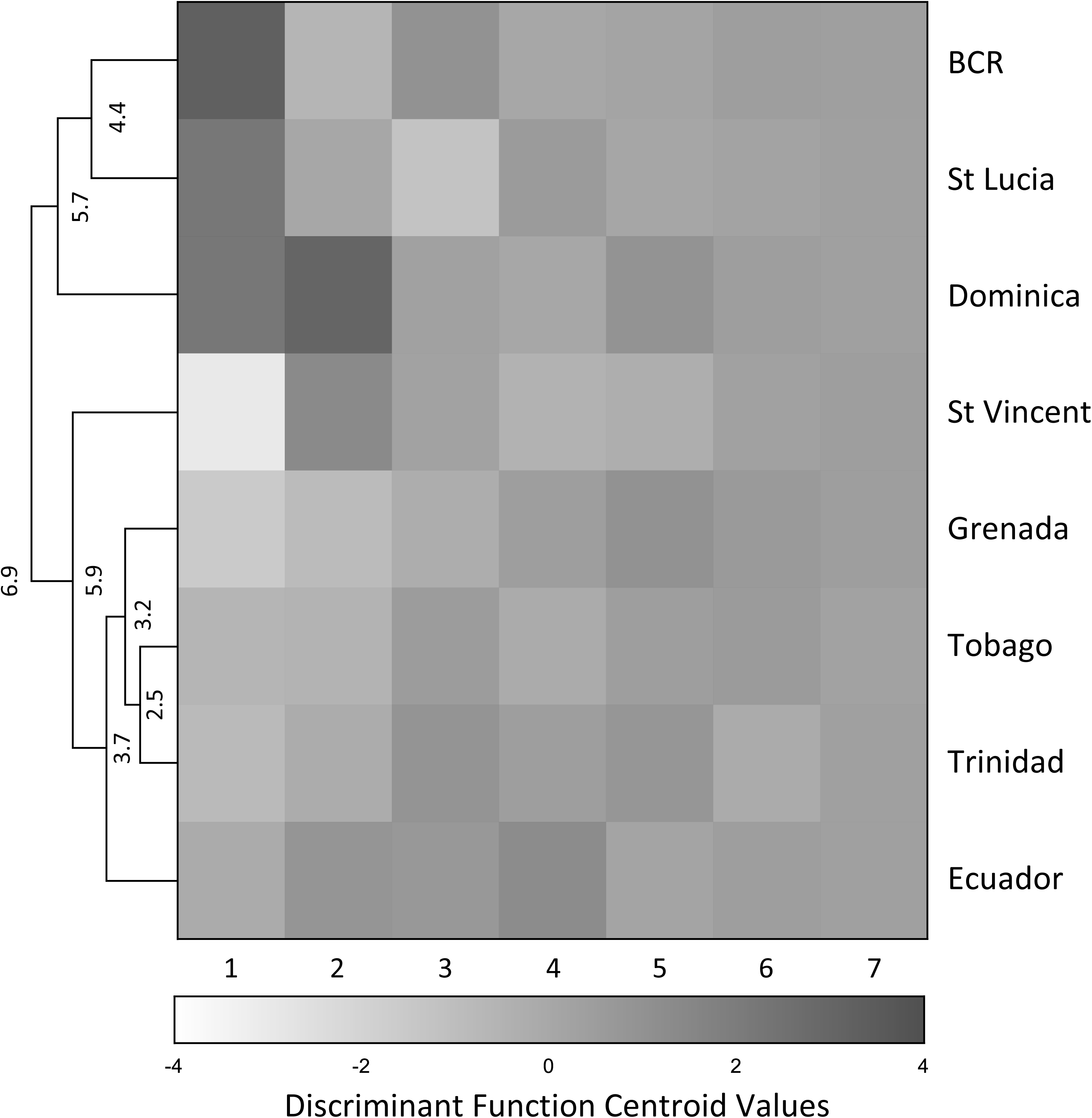
Bivariate plots of discriminant scores for the songs of each population on the first three discriminant functions (a-c) and a heat map and clustered dendrogram (d) summarizing the relationships among populations based on the multivariate distances between them. The first panel (a) shows some separation of BCR, Dominica, St Vincent and to a lesser extent Grenada on the first two discriminant functions, while the second panel (b) shows that the third discriminant function helps to separate St Lucia and it also highlights (with large elipses) the distinct clustering of BCR, St Lucia and Dominica seaparately from Grenada, Trinidad, Tobago, and St Vincent. The third panel (c) shows no further separation of the populations. The fourth panel (d) is a heat map of population centroid values on each of the seven functions from discriminant analysis and a clustered dendrogram that groups the populations based on similarity and difference in the cumulative euclidean distance between their centroid values across all seven functions using the group averaging clustering method. Values at the nodes of the dendrogram represent the average distance between the branches they connect, expressed in units defined by the discriminant space: the absolute values therefore have no wider meaning but they can be interepreted in relative terms, such that the relative differences in the magnitude of the node values capture the relative magnitude of the song differences between the populations. So, for example, the difference in songs linking St Lucia and BCR is greater than the difference in songs that links the entire cluster of populations of Grenada, Tobago, Trinidad, and Ecuador.

Classification success from discriminant analysis using all seven functions is shown in Table 7 which confirms and more clearly quantifies the patterns of overlap versus separation among the populations observed in the discriminant score plots. Overall, 65.9% of songs were correctly classified, which represents a significant degree of overall discrimination among populations and a very considerable improvement on random (chance) classification which would be approximately 12.5% for this sample. There was also considerable variation in levels of classification success for the different populations. Of the island populations, St Vincent (85.7% correct) and Dominica (80.0%) were correctly classified at the highest levels, followed by St Lucia (65.6%) and Grenada (60.0%), while Tobago (33.3%) and Trinidad (25.8%) showed the lowest levels of successful classification. Both mainland populations were classified at relatively high levels (BCR: 77.1%; Ecuador 79.7%).

**Table 7.** Classification results from discriminant analysis.

| Population | N | BCR | Dominica | StLucia | StVincent | Grenada | Tobago | Trinidad | Ecuador | % correct | Certainty Own* |
| --- | --- | --- | --- | --- | --- | --- | --- | --- | --- | --- | --- |
| BCR | 48 | **37** | 2 | 4 | 0 | 1 | 2 | 0 | 2 | 77.1% | 0.72 |
| Dominica | 30 | 2 | **24** | 0 | 0 | 0 | 0 | 0 | 4 | 80.0% | 0.76 |
| StLucia | 32 | 4 | 2 | **21** | 0 | 1 | 1 | 1 | 2 | 65.6% | 0.62 |
| StVincent | 28 | 0 | 0 | 0 | **24** | 1 | 1 | 1 | 1 | 85.7% | 0.77 |
| Grenada | 30 | 0 | 0 | 1 | 3 | **18** | 2 | 2 | 4 | 60.0% | 0.47 |
| Tobago | 30 | 1 | 0 | 0 | 1 | 6 | **10** | 5 | 7 | 33.3% | 0.33 |
| Trinidad | 31 | 0 | 0 | 0 | 2 | 6 | 4 | **8** | 11 | 25.8% | 0.31 |
| Ecuador | 64 | 2 | 0 | 1 | 1 | 1 | 4 | 4 | **51** | 79.7% | 0.63 |
| **Totals:** | **293** |  |  |  |  |  |  |  |  | **65.9%** |  |
| #Misclassified to: |  | 6 | 4 | 6 | 7 | 16 | 14 | 13 | 31 |  |  |
| %Misclassified to: |  | 6.7% | 4.3% | 6.7% | 7.3% | 18.2% | 17.5% | 16.9% | 35.6% |  |  |

\* Certainty own represents the probability that a song from a given population will be assigned to itself, as opposed to a different population, averaged across all songs for that population. Average values for each population express the confidence of correct self-assignment.
Misclassified totals and % values for each population are based on the total of the misclassified cases that **could possibly be assigned to them**, which therefore excludes their own misclassified songs that were assigned to other populations. It is an additional measure of distinctiveness that captures how likely a given population is to be mistaken for other populations.

Table 7 also tabulates a “Certainty Own” statistic which repesents the probability that a song from a given population will be assigned to itself, as opposed to a different population, averaged across all songs for that population. The statistic is similar but not equivalent to % correct because any given song can be assigned correctly (or incorrectly) with varying certainty based on the relative probabilities that it could belong to one or more of the other populations. So the statistic shows, for example, that songs from Dominica and St. Vincent are not only correctly classified at high levels (80-85%) but also with comparatively high certainty (>75%), indicating that there is little possibility that they belong to another population, which further underscores the relative distinctiveness of these two populations from all the others. In contrast, songs from Grenada are classified moderately well (60.0%) but with relatively low certainty (47%) confirming their broader similarity to other populations.

Finally, Table 7 also shows that misclassified cases were not randomly distributed among the populations. The small number of cases for Dominica that were misclassified were not assigned to any of the other island populations but rather only to either BCR or Ecuador. A similar bias held for St Lucia with more misclassified cases assigned to either BCR or Dominica and only one or two assigned to any of the other populations. The remaining populations showed a reciprocal bias with misclassified cases from Grenada, Tobago and Trinidad all being assigned either to one another or to Ecuador but not to St Lucia, Dominica or BCR; and misclassified cases from Ecuador were assigned primarily to Trinidad and Tobago. St Vincent had very few misclassified cases, but they were also all assigned to either Grenada, Tobago, Trinidad or Ecuador and not to St Lucia, Dominica or BCR.

These patterns in misclassification are summarized in the final rows of Table 7, which tally the number and percentage of all misclassified cases that were assigned to each of the different populations. This serves as another measure of the relative distinctiveness of each population by capturing how likely a given population is to be mistaken for other populations. These tallies further confirm the distinctiveness of Dominica, St Lucia and St Vincent: not only were these three populations successfully classified at the highest levels, they were also not often mistaken for any of the other populations. By comparison Grenada, Tobago and Trinidad were more often mistaken for another population in addition to being successfully classified at lower levels.

The overall patterns of similarity and difference among the populations are distilled in the final panel in Figure 6 in the form of a heat map and clustered dendrogram. The heat map plots the centroid values for each population on each of the seven discriminant functions represented by the variable grey-scale shading of each box. Boxes that are more similarly shaded have similar centroid values (i.e., are closer to each other in multivariate space) and the grey-scale patterns here simply replicate what is graphically evident also in the discriminant score plots in the preceding three panels. The clustered dendrogram distills the overall results pattern by grouping populations based on the cumulative similarity or difference between their centroid values across all seven functions. The dendrogram groups BCR, St Lucia and Dominica together and separates them from the other populations as a distinct cluster. It then groups Tobago with Trinidad, followed by Grenada, and then Ecuador, with St Vincent as the final and more distant member of the second cluster.

## DISCUSSION

### Summary Findings

In their general structure, organization and delivery the song patterns of island populations of House Wren in the Lesser Antilles are all very similar and similar also to continental forms of northern and southern House Wren (Kroodsma 1977; Tubaro 1990; Rendall and Kaluthota 2013; Sosa-López and Mennill 2014a, b; dos Santos et al. 2016; DiSciullo et al. 2023) and notably also to some of the other island populations in Mexico whose song has been comprehensively described (Sosa-López and Mennill 2014c). Thus, they all show the same basic organizational template, with songs structured in two discrete sections, an Introduction section of lower amplitude often noisy elements, and a Main section of louder, tonal and frequency modulated elements; they all show the same pattern of song type delivery where bouts of songs involve serial repetition of the same song type many times before switching and then successive song types tend to differ only minimally by the addition, deletion, substitution or movement of a single syllable type at a time; and they all also show a similar capacity to deviate from both of these patterns. Thus, songs sometimes omit one or other of the Introduction or Main sections; at other times they involve concatenating a series of them together into song ‘doublets’ or ‘triplets’. Similarly, sequences of songs are often very repetitious, but song sequences at other times can be far more variable and involve rapid switching between different song types with more wholesale changes in the note and syllable contents of successive songs. The latter sorts of deviation from the basic structural and organizational templates may be developed and used to a greater or lesser extent in some of the populations, but in these higher-level characteristics, the song patterns of all populations are generally more similar than they are different.

There are however diagnosable differences in the more detailed features of individual songs. Thus, songs in Dominica are markedly longer in overall duration, with the longest Main section, the longest intervals between elements, and thus also one of the slowest rates of element production. They also entail a preponderance of protracted broad-band frequency sweeps that yield the highest average center frequency and markedly higher averages for the fundamental frequency (F_0_) and bandwidth of elements in the Main section. Collectively, the impression is of a kind of relaxed, or largo, delivery style with songs produced at a slower tempo with widely spaced, broadband trilled elements. These particular findings agree well with the song analysis by Sosa-López and Mennill (2014a). Although Dominica was the only island population from the Lesser Antilles that was represented in that broad-scale study of house wrens, it was described to have the most distinctive song with specific distinguishing characteristics very much in agreement with those identified here.

Songs from St Vincent are also notably distinctive. They have the highest average center frequency for the Introduction section and the shortest Main section, with the fewest number of elements of markedly lowest bandwidth, frequency modulation, and average entropy (the latter measure capturing the relatively pure tone nature of elements in the Main section). Subjectively, the impression is of a relatively high frequency and often shrill, or even discordant, introductory section paired with a short simple Main section with comparatively few steretoytped, narrowband elements that are slightly offset in frequency creating a high-low, sing-song quality.

Songs from StLucia are – subjectively – more typically House Wren but may be discriminably different in having a relatively short Introduction section with fewer and more tonal elements (with lowest average entropy and highest signal-to-noise ratio) combined with a proportionally longer Main section containing more elements. The impression is virtually the opposite of that for Dominica, entailing a more rapid fire – staccato – delivery style with a relatively long and complex Main section.

Songs from Grenada are potentially distinctive in having the shortest Introduction section with the fewest number of elements, contributing to the shortest overall song duration where the Main section represents a greater proportion of the song relative to other populations. Otherwise, they bear the most resemblance to canonical House Wren song, being intermediate in almost every dimension and most similar to Tobago, Trinidad and Ecuador.

Songs from Tobago and Trinidad are not exceptional on any of the measured acoustic features, although Main section elements have comparatively low bandwidth, and the two populations are similar to each other in this and most other features.

Taken together, the picture that emerges for song of the various island populations of the Lesser Antilles is of a broadly conserved and over-arching organizational template common to all house wrens, with population-specific differences arising in the more detailed features of song structure, where Dominica and St Vincent are most distinctive, followed by St Lucia and Grenada, with Trinidad and Tobago minimally distinctive from mainland populations.

### Taxonomic Implications: How Many Species?

Overall, these findings provide some important empirical support for the recent taxonomic reclassification that elevated to species status the four most distant islands of the Lesser Antilles, namely Dominica, St Lucia, St Vincent and Grenada (Chesser et al. 2024). In discriminant analysis of detailed song features, these four populations were discriminated at levels significantly above chance and substantially higher than levels of success for either Trinidad or Tobago. Of course, statistical discrimination is not by itself definitive proof of species differentiation. Nevertheless, it does have an important bearing on the issue because, as reviewed in the Introduction, each of the major forms of data available previously and used in support of the taxonomic splits were either fragmentary or incomplete in their sampling of the different island populations, or equivocal in the patterns manifest. In such situations, some weight in taxonomic exercises is often attached to song given its role in mate recognition and choice (Remsen 2005). Hence, the findings here, based on a comprehensive sampling and analysis of song across all of the islands, represents an important contribution to the taxonomy of this group and provides support for most of the splits. The cases are strongest for Dominica and St Vincent whose songs are most distinctive, and to a somewhat lesser extent also for St Lucia. The case for species status – at least based on song – is arguably weakest for Grenada which was successfully classified at the lowest levels of the four islands in question (though considerably better than either Tobago or Trinidad) and was also more frequently mistaken for one of the other populations.

Importantly also, the discriminant analysis specifically confirmed the similarity of songs between Trinidad and Tobago, and the similiarity of both to songs from mainland South America, as represented by populations in Ecuador. This similarity in song patterns among all three populations may be evidence of some continuing contact (and gene flow) among them which would not be unexpected given the proximity of Trinidad and Tobago to each other and to mainland South America. It is therefore consistent with, and further supports, the recent taxonomic exercise which identifies them all as subspecies of the mainland south American form (*T. musculus*; Chesser et al. 2024)

### Evolutionary Implications

An additional novel result of this study, with important evolutionary implications, is the summary finding from discriminant analysis of two distinct clusters of island populations: one cluster that groups Dominica and St Lucia (the two most northern islands in the chain) as distinct from the others and linked most closely to BCR as the most similar mainland population; and another cluster that groups Tobago, Trinidad and Grenada (the three most southern islands) linked most closely to Ecuador, with St Vincent (in the middle of the island chain) an outlier to this second cluster (Figure 6). This pattern provides some support for, and also extends, the intriguing but tentative hypothesis from the genetic study by Klicka et al. (2023) that there might be different geographic sources for the various islands in the lesser Antilles, specifically that birds in Dominica might originate from Central America, while birds on the other islands likely have a south American origin. This hypothesis was tentative given the limitations of mtDNA data (reviewed in Winker 2021) and the absence of complementary nuclear DNA results for the Lesser Antilles populations; and the possibility of a central American source was suggested only for Dominica because Klicka et al. (2023) did not have any genetic samples for St Lucia. So the results here for song data are productive in that they align with the broader point of the Klicka et al. hypothesis that the island populations of the Lesser Antilles might have different continental sources, and the results here specifically extend that proposal to include St Lucia in those with a possible central American origin.

This is important because, if true, the link of both Dominica and St Lucia – at the north end of the island chain – to a mainland source from Central America, distinct from the other islands with a likely source from South America, makes all the more interesting the patterns of distinctive variation in other traits across the islands. For example, the most noticeable (and commonly noted) pattern of island variation is in plumage, where the four islands recently raised to species status are all different in plumage from mainland forms but themselves actually sort into two pairs of very similarly colored morphs: one pair links Dominica and Grenada which are at the northern and southern extremes of this group of four islands; and the other pair links St Lucia and St Vincent which are in the middle (see Figure 1 and the Introduction section for more details). Hence, if Dominica and St Lucia (at the north end) share a central American source, while St Vincent and Grenada (at the south end) share a south American source, then the leap-frog pattern of plumage similarity and difference among them would require two independent instances of convergent plumage divergence: in other words, two separate instances where the divergence in plumage within each of the two pairs resulted in convergence on very similar plumage patterns across them. It is not immediately obvious what set of factors might naturally explain such convergent divergence but it does not map naturally onto the song patterns among them either: Dominica and Grenada are similar in plumage but not in song; St Lucia and St Vincent are likewise similar in plumage but not in song; while Grenada, Trinidad and Tobago are most similar in song but not in plumage.

### Female Song

Adding to this already interesting character mix is the pattern of female song across the islands, in so far it is known, which is very poorly. Although not a focus of study, my own observations (and other informal reports) confirm the presence of female song on all of the islands except St Lucia and St Vincent. This is curious because female song is common in tropical songbird species generally, and it is also known, though possibly variably used, in continental populations of both northern and southern House Wren (Johnson and Kermott 1990; Krieg and Getty 2016; Keck et al. 2025; reviewed in Fernandez et al. 2024; Johnson 2024). Where it occurs among house wrens in the Lesser Antilles, female song is similar to continental counterparts and often takes the form of a short sequence of relatively harsh unstructured notes, similar to some introductory notes of male song and some common non-song vocalizations. It can, however, also include more tonal, frequency modulated notes similar to some of the more structured elements of the Main section of male song (for an example from Dominica, see Figure 1 in Johnson et al. 2025). And female song can at times be highly synchronized to the male’s song, appended immediately to the end of his song almost as an accentuation or echo of it; at other times it overlaps male song and can be difficult to distinguish from it; and yet other times it can lead or occur completely independently of male song.

This is a very cursory and preliminary characterization of female song, unfortunately, and the matter deserves much more detailed study because, while not historically a focus of research to the same extent as male song, very recent comprehensive comparative studies and evolutionary path analysis have shown that female song is both more widespread across songbird taxa than previously appreciated and likely also represents the ancestral state for many of them (Odom et al. 2014; see also Odom and Benedict 2018). This work further shows that, across tropical and temperate zone taxa, the absence of female song likely reflects evolutionary loss associated with a shift away from year-round territoriality and biparental care (Odom et al. 2025). Hence, if female song is truly absent in the Lesser Antilles islands of St Lucia and St Vincent, that have putatively different continental sources, it would not only represent an additional instance of convergent evolutionary divergence – the independent loss of song in two island populations with different original sources – but it might also point to associated evolutionary changes in other core behavioral and life-history traits. Further, if effective mate choice and pairing involves coordinating song contributions between prospective male and female partners, then the absence of female song in St Lucia and St Vincent could be an important taxonomic character as an effective premating isolating mechanism.

These latter possibilities remain speculative but there currently appear to be multiple possible instances of character divergence and convergence among plumage, male song and possibly also female song (and potentially other behavioral and life-history traits) across the islands of the Lesser Antilles. Collectively, this intriguing mix of characters therefore stands as a strong impetus to further detailed study to bring additional light to the evolutionary relationships among these island populations of house wrens and to the historical processes that have yielded the trait mosaic they now manifest.

### Summary and Future Research Directions

1. In general structure, organization and delivery the song of males on all of the islands of the Lesser Antilles is similar and follows patterns common to house wrens broadly, but songs of the different islands are discriminably different in their more detailed features.
2. The latter differences in song among the island populations are consistent with most of the recently recognized species splits, with the possible exception of Grenada where song is the least distinctive and comparatively closely aligned with that of Trinidad, Tobago and Ecuador. Birds in Grenada are distinct from continental forms (and Trinidad and Tobago) in plumage but this difference is of uncertain classification value given the environmental plasticity of plumage (and size) generally and the fact that birds in Grenada are most similar in plumage to birds at the other end of the island chain in Dominica which genetic and song analysis both link to a different geographic route of incursion.
3. Additional genetic research, especially including nuclear DNA, would help to resolve the relationships among island populations and clarify their potential central versus south American origins, which would in turn bring additional clarity to the character mosaic manifest among them that currently implies multiple independent instances of evolutionary convergence and divergence.
4. Female song should be a particular focus of future research to confirm or revise its presence on each of the islands and to more fully characterize its features and possible functions, including potential coordination with male song.

## Acknowledgements

I am grateful to Karen Rendall for field assistance in 2015; to Rochelle Bellemare and Rupert Radix for logistical support at Simla Field Research Station and the Asa Wright Nature Center in Trinidad; to ProAves and OTS, respectively, for visits to Copalinga Reserve in Ecuador and the Las Cruces Biological Research Station in Costa Rica; and to Theresa Burg for several helpful discussions of the topic and comments on the manuscript. Research funding was provided from the Natural Sciences and Engineering Research Council of Canada (NSERC) through the Government of Canada’s Tri-Agency research funding program, the University of Lethbridge, and the University of New Brunswick.

Research adhered to guidelines of the Canadian Council on Animal Care and was conducted in accordance with approved animal welfare protocols from the University of Lethbridge (AWC#1023; AWC#1429) and the University of New Brunswick (ACC#18041, ACC#19055). I report no conflicts of interest and am responsible for all aspects of the research reported. In accordance with institutional and Tri-Agency policy, research data will be deposited in the UNB-DATAVERSE.

## LITERATURE CITED

1. Alström, Per & Ranft, Richard. (2003). The use of sounds in avian systematics and the importance of bird sound archives. Bull. B.O.C.. 123.

2. American Ornithologist’s Union (1983). Check-list of North American Birds, 6th ed. Am, Ornithol. Union, Washington, D.C.

3. American Ornithologist’s Union (1998). Check-list of North American Birds, 7th ed. Am, Ornithol. Union, Washington, D.C.

4. Bailey, F.M. (1902). Handbook of Birds of the Western United States. Houghton, Mifflin and Co., Cambridge, Massachusetts.

5. Barlow, J.C. (1978). Another colony of the Guadeloupe house wren. The Wilson Bulletin 90 (4): 6356–637

6. Boersma, P. , and D. Weenink (2021). PRAAT Software, version 6.1.40

7. Brewer, D. (2001). Wrens, Dippers and Thrashers. Yale University Press.

8. Brumfield, R. T., and A. P. Capparella. (1996). Genetic differentiation and taxonomy in the house wren species group. Condor 98:547–556.

9. Center for Conservation Bioacoustics (2019). Raven Pro: Interactive Sound Analysis Software (Version 1.6.1) Ithaca, NY: The Cornell Lab of Ornithology

10. Chesser, R. T., S. M. Billerman, K. J. Burns, C. Cicero, J. L. Dunn, B. E. Hernández-Baños, R. A. Jiménez, O. Johnson, A. W. Kratter, N. A. Mason, P. C. Rasmussen, and J. V. Remsen, Jr. (2024). Sixty-fifth Supplement to the American Ornithological Society’s Check-list of North American Birds. Ornithology 141: 1–20. 10.1093/ornithology/ukae019

11. Clegg, S. M., and I. P. F. Owens (2002). The ‘island rule’ in birds: Medium body size and its ecological explanation. Proceedings of the Royal Society of London B 269: 1359–1365.53

12. Cox, G. W. and R. E. Ricklefs (1977). Species diversity and ecological release in Caribbean land bird faunas. Oikos 28(2): 113–122.

13. Cyr, M.E., K. Wetten, M.H. Warrington, and N. Koper (2021). Variation in song structure of house wrens living in urban and rural areas in a Caribbean small island developing state. Bioacoustics 30 (5): 594–607. DOI: 10.1080/09524622.2020.1835538

14. Deslandes, V., L. R. R. Faria, M. E. Borges and M. R. Pie. (2014). The structure of an avian syllable syntax network. Behavioural Processes 106:53–59.

15. DiSciullo, R.A., S. K. Sakaluk, and C. F. Thompson (2023). Song structure of male Northern House Wrens and patterns of song production and delivery across the nesting cycle. J Ornithology 10.1007/s10336-023-02098-0

16. DiSciullo, R.A., A.M. Forsman, R. R. Fitak, J. Hunt, P. Nietlisbach, C. F. Thompson, and S. K. Sakaluk (2024). Male song structure predicts offspring recruitment to the breeding population in a migratory bird. Evolution 78(6): 1054–1066.

17. Dickinson, E. C., and L. Christidis, Editors (2014). The Howard and Moore Complete Checklist of the Birds of the World. Volume 2. 4th edition. Aves Press, Eastbourne, UK. ISBN 978-0-9568611-2-2.

18. Fernández, G.J., M.E. Carro, and L. S. Johnson (2024). Southern House Wren (*Troglodytes musculus*), version 1.0. In Birds of the World (R. Juárez, B. K. Keeney, and S. M. Billerman, Editors). Cornell Lab of Ornithology, Ithaca, NY, USA. 10.2173/bow.houwre4.01

19. Gilardi, J. D., and C. L. John (1998). Conservation of the St. Lucia House Wren *Troglodytes aedon mesoleucus*: Distribution, abundance and breeding biology. *Dodo*, Jersey Wildlife Preservation Trust 34:91–102.

20. Grant, P. R. (1965). The adaptive significance of some size trends in island birds. Evolution 19(3): 355–367.

21. Grant, B. R., and P. R. Grant (1996). Cultural inheritance of song and its role in the evolution of Darwin’s finches. Evolution 50(6): 2471–2487

22. Huber, S., and J. Podos (2006). Beak morphology and song features covery in a population of Darwin’s finshes (*Geospiza fortis*). Biological Journal of the Linnean Society 88: 489–498.

23. Hunt, P. D., and D. J. Flaspohler (2020). Yellow-rumped Warbler (*Setophaga coronata*), version 1.0. In Birds of the World (P. G. Rodewald, Editor). Cornell Lab of Ornithology, Ithaca, NY, USA. 10.2173/bow.yerwar.01

24. Irwin, D.E. (2000). Song variation in an avian ring species. Evolution 54: 998–1010.

25. Johnson, L. S. (2024). Northern House Wren (*Troglodytes aedon*), version 1.1. In Birds of the World (B. K. Keeney, A. F. Poole, M. G. Smith, and S. M. Billerman, Editors). Cornell Lab of Ornithology, Ithaca, NY, USA. 10.2173/bow.houwre.01.1

26. Johnson, L. S., and L. H. Kermott (1990). The structure and context of female song in a north-temperate population of House Wrens. Journal of Field Ornithology 61:273–284.

27. Johnson, L. S., and L. H. Kermott (1991). The functions of song in male House Wrens (*Troglodytes aedon*). Behaviour 116: 190–209.

28. Johnson, L. S., and W. A. Searcy. (1996). Female attraction to male song in House Wrens (*Troglodytes aedon*). Behaviour 133:357–366.

29. Johnson, L. S., R. Juárez, and D. Rendall (2025). Kalinago Wren (*Troglodytes martinicensis*), version 1.1. In Birds of the World (B. K. Keeney, S. M. Billerman, J. Gerbracht, and A. J. Spencer, Editors). Cornell Lab of Ornithology, Ithaca, NY, USA 10.2173/bow.cinwre1.01.1

30. Juárez, R., L. S. Johnson, and D. Rendall (2025). St. Lucia Wren (*Troglodytes mesoleucus*), version 1.1. In Birds of the World (B. K. Keeney, S. M. Billerman, and A. J. Spencer, Editors). Cornell Lab of Ornithology, Ithaca, NY, USA. 10.2173/bow.houwre8.01.1

31. Kaluthota, C., B. E. Brinkman, E. B. dos Santos, and D. Rendall (2016). Transcontinental latitudinal variation in song performance and complexity in house wrens (*Troglodytes aedon*). Proceedings of the Royal Society of London B. 283:20152765.

32. Kaluthota, C.D, Logue, D.M., and Rendall, D. (2020). Conventional and network analyses of song organization and complexity in northern House Wrens (*Troglodytes aedon parkmanii*). Journal of Field Ornithology. 10.1111/jofo.12347

33. Keck, SM., A. Barnhill, S. Q. Smeele, M. H. Hsu, A. Dolson-Fazio, C. Bergler, G. F. Ball, C. A. Krieg and K. J. Odom (2025) Deep machine learning allows classification of male and female songbird songs in a species with highly variable repertoires Behaviour DOI: 10.1163/1568539X-bja10324

34. Klecka, W.R. (1980). Discriminant Analysis Series: Quantitative Applications in the Social Sciences (Sage, London).

35. Klicka, J., K. Epperly, B. T. Smith, G. M. Spellman, J. A. Chaves, P. Escalante, C. C. Witt, R. Canales-del-Castillo, and R. M. Zink (2023). Lineage diversity in a widely distributed New World passerine bird, the House Wren. Ornithology 140:1–13. 10.1093/ornithology/ukad018

36. Krieg, C.A, and T. Getty (2016). Not just for males: females use song against male and female rivals in a temperate zone songbird. Animal Behaviour 113: 39–47

37. Kroodsma, D.E. (1977). Correlates of song organization among North American wrens. American Naturalist 109: 995–1008.

38. Kroodsma, D. E., and D. Brewer (2005). Family *Troglodytidae* (wrens). In Handbook of the Birds of the World. Volume 10: Cuckoo Shrikes to Thrushes (J. del Hoyo, A. Elliott, and D. A. Christie, Editors). Lynx Edicions, Barcelona, Spain. pp. 356–447.

39. Linsdale, J. M. (1928). Variations in the Fox Sparrow (*Passerella iliaca*) with reference to natural history and osteology. University of California Publications in Zoology 30:251–392.

40. Valderrama, S., J. Parra, and D.J. Mennill (2007). Species differences in the songs of the critically endangered Niceforo’s Wren and the related Rufous-and-white Wren. Condor 109(4): 871–878.

41. NCSS 2023 Statistical Software (2023). NCSS, LLC. Kaysville, Utah, USA, ncss.com/software/ncss Oberholser, H. C. (1904). A review of the wrens of the genus *Troglodytes*. Proceedings of the United States National Museum 27:197–210.

42. Odom, K.J., and L. Benedict (2018). A call to document female bird songs: applications for diverse fields. Auk 135: 314–325. DOI:10.5751

43. Odom, K.J., Hall, M.L., Riebel, K., Omland, K.E., and Langmore, N.E. (2014). Female song is widespread and ancestral in songbirds. Nature Communications 5: 3379. DOI:10.1038/ncomms4379.

44. Odom, KJ, M Araya-Salas, L Benedict, K Lim, J Dale, WH Webb, C Sheard, J.A. Tobias, G. F. Ball, M.L. Hall, N.E. Langmore, M.S. Webster, and K. Riebel (2025). Global incidence of female birdsong is predicted by territoriality and biparental care in songbirds Nature Communications 16 (1), 6157

45. Podos, J., J.A. Southall, and M.R. Rossi-Santos (2004). Vocal mechanics in Darwin’s finches: correlation of beak gape and song frequency Journal Experimental Biology (2004) 207 (4): 607–619.

46. Podos, J. (2010). Acoustics discrimination of sympatric morphs in Darwin’s finches: a behavioral mechanism for assorative mating? Phil Trans Royal Society, B 365: 1031–1039.

47. Remsen, J.V. (1984). High incidence of" leapfrog" pattern of geographic variation in Andean birds: implications for the speciation process. Science, 224 (4645): 171–173.

48. Remsen, J.V. (2005). Pattern, process and rigor meet classification. The Auk, 122(2): 403–413

49. Rendall, D., and C. D. Kaluthota. (2013). Song organization and variability in Northern House Wrens (*Troglodytes aedon parkmanii*) in western Canada. Auk 130 (4):617–628.

50. dos Santos, E.B., P.E. Llambías, and D. Rendall (2016). The structure and organization of song in Southern House Wrens (*Troglodytes aedon chilensis*). Journal of Ornithology. 157(1): 289–301.

51. dos Santos, E.B., Llambías, P., and D. Rendall (2018). Male song diversity and its relation to breeding success in Southern House Wrens (*Troglodytes aedon chilensis*). Journal of Avian Biology DOI: 10.1111/jav.01606

52. Sosa-López, J. R., and D. J. Mennill (2014a). Continent-wide patterns of divergence in acoustic and morphological traits in the House Wren species complex. The Auk 131(1):41–54.

53. Sosa-López, J. R., and D. J. Mennill (2014b). The vocal behavior of the Brown-Throated Wren (*Troglodytes brunneicollis*): song structure, repertoires, sharing, syntax, and diel variation. J Ornithology 115: 435–446.

54. Sosa-López, J. R., and D. J. Mennill (2014c). Vocal behavior of the island-endemic Cozumel Wren (*Troglodytes aedon beani*): song structure, repertoires, and song sharing. J Ornithology 115: 337–346.

55. Sosa-López, J. R., J. E. Martínez Gómez, and D. J. Mennill (2016). Divergence in mating signals correlates with genetic distance and behavioural responses to playback. Journal of Evolutionary Biology 29:306–318. 10.1111/jeb.12782

56. Sukumaran, J., and L. Knowles (2021). Multispecies coalescent delimits structure, not species. Proc Nat Acad Sci 114: 1607–1612. www.pnas.org/cgi/doi/10.1073/pnas.1607921114

57. Swarth, H. W. (1920). Revision of the avian genus *Passerella* with special reference to the distribution and migration of the races in California. University of California Publications in Zoology 21:75–224.

58. Tabachnick, B.G., and L.S. Fiddell (2007). Using Multivariate Statistics, 5th edition. Pearson: New York.

59. Toews, D.P.L, and D.E. Irwin (2008). Cryptic speciation in a Holarctic passerine revealed by genetic and bioacoustic analyses. Molecular Ecology 17: 2691–2705.

60. Tubaro, P. L. (1990). Song description of the House Wren (*Troglodytes aedon*) in two populations of eastern Argentina, and some indirect evidences of imitative vocal learning. Hornero 13:111–16.

61. Weckstein, J. D., D. E. Kroodsma, and R. C. Faucett (2020). Fox Sparrow (*Passerella iliaca*), version 1.0. In Birds of the World (A. F. Poole and F. B. Gill, Editors). Cornell Lab of Ornithology, Ithaca, NY, USA. 10.2173/bow.foxspa.01

62. Wetten, K. N. (2021). Morphological divergence in the House Wren (Troglodytes aedon) species complex: A study of island populations with a focus on the Grenada House Wren (T. a. grenadensis). M.Sc. thesis, University of Manitoba, Canada.

63. Wiebe, K. L., and W. S. Moore (2024). Northern Flicker (*Colaptes auratus*), version 2.1. In Birds of the World (P. G. Rodewald and B. K. Keeney, Editors). Cornell Lab of Ornithology, Ithaca, NY, USA. 10.2173/bow.norfli.02.1

64. Winker, K. (2021). An overview of speciation and species limints in birds. Ornithology 138: 1–27. DOI: 10.1093/ornithology/ukab006

65. Young, B.E. (1994). The effects of food, nest predation and weather on the timing of breeding in tropical house wrens. Condor 96: 341–353

66. Zink, R. M. (1994). The geography of mitochondrial DNA variation, population structure, hybridization, and species limits in the Fox Sparrow (*Passerella iliaca*). Evolution 48:96–111.

67. Zink, R.M. (2008). Microsatellite and mitochondrial DNA variation in the Fox Sparrow. The Condor 110 (3): 482–492.

68. Zink, R. M., and R. C. Blackwell (1996). Patterns of allozyme, mitochondrial DNA, and morphometric variation in four sparrow genera. The Auk 113(1):59–67.

69. Zink, R. M., and J. D. Weckstein (2003). Recent evolutionary history of the Fox Sparrows (Genus: *Passerella*). The Auk 120(2):522–527.

